# Sludge liquefaction and stratification enable ultra-high digestion rate and resource recovery for waste activated sludge treatment

**DOI:** 10.1101/2025.04.27.650825

**Authors:** Shanquan Wang, Qihong Lu, Zhiwei Liang, Chen Wang, Jiangjian Shi, Boyu Jia, Yuhan Liang, Daqian Jiang, Dawei Liang, Yang Zhang, Jan Dolfing, Peng Wang, Lei Wang, Rongliang Qiu

**Author notes:** These authors contributed equally (Shanquan Wang, Qihong Lu, Zhiwei Liang). Corresponding authors: Shanquan Wang Address: School of Environmental Science and Engineering, Sun Yat-Sen University, Guangzhou, China 510275 Email address, Rongliang Qiu Address: Guangdong Laboratory for Lingnan Modern Agriculture, Guangdong Provincial Key Laboratory of Agricultural & Rural Pollution Abatement and Environmental Safety, College of Natural Resources and Environment, South China Agricultural University, Guangzhou, China 510642; School of Environmental Science and Engineering, Sun Yat-sen University, Guangzhou, China 510006.

## Abstract

Most of the energy, nitrogen/N and phosphorus/P entering wastewater treatment plants (WWTPs) accumulates in waste activated sludge (WAS). While these resources are theoretically recoverable through anaerobic digestion (AD), conventional sludge AD faces long-standing challenges of low organic loading rate and limited N/P recovery. Here, building on an innovative sludge liquefaction and stratification technique, we present a novel process entitled SPREAD that boasts 10 times higher organic loading rates and recovers 5 folds of N/P resources, relative to conventional sludge AD. We develop an anaerobic digestion database (ADDB) for multi-omics analyses of digestion microbiomes, and pinpoint the mechanism underlying the exceptionally high organic loading rate and remarkable N/P recovery in SPREAD. Importantly, we identify a negative correlation between the Gibbs free energy (ΔG) and the relative abundance of methanogens, enabling identification of bottleneck steps for stimulation and augmentation of methanogenic digestion. Our study provides a game-changing technology for the treatment and resource recovery from WAS and opens a new avenue for sustainable management and carbon/energy-neutrality of WWTPs.

## Main

Each year, approximately 3.80 × 10^11^ m^3^ of municipal wastewater is generated globally, so wastewater treatment plants (WWTPs) can be resource factories to recover valuable chemical and energy resources^1,2^. In this wastewater-resource factory scenario, WWTPs could generate 6.5 × 10^9^ kWh of electrical energy, and recover 1.9 × 10^7^ tonnes of N and 3.8 × 10^6^ tonnes of P each year, rather than consuming 3% of global electricity by current WWTPs^1–3^. Moreover, around 50% reducing equivalents, together with 50% N and 90% P resources, flow to the activated sludge basin during wastewater treatment, resulting in concentration of N/P and energy resources in waste activated sludge (WAS)^4,5^. Thus far, anaerobic digestion (AD) is the most widely used approach to reduce and stabilize WAS via organics-to-biogas conversion, which offset 20-30% of the energy and greenhouse-gas costs of the activated sludge process^1,6^. In addition, a small portion (<10%) of sludge N/P resources can be recovered from the digestion supernatant via air stripping and struvite precipitation^7,8^. However, application of sludge AD is severely hampered by the following issues: (i) low organic loading rate and consequent large footprint due to long sludge retention time in AD reactors; the organic loading rates of conventional sludge AD generally range from 0.5 to 1 kg COD m^-^³ day^-1^, with 20-30 days of sludge retention time^9,10^; (ii) low solid-solid mass transfer coefficient in sludge AD; a bottleneck in sludge digestion is the organic mass transfer between sticky floc-like sludge and digesting microorganisms, which restricts substrate diffusion and microbial accessibility^11,12^; (iii) ammonia inhibition; sludge AD involves hydrolysis, acidogenesis, acetogenesis and methanogenesis steps, all of which functional microorganisms are sensitive to high concentrations of sludge-protein-derived ammonia, particularly the archaeal methanogens^13–15^. For instance, 560 mg/L free ammonia can reduce methanogenic activities by 50%^14^. Sludge pretreatment can improve the overall organics-to-biogas conversion efficiency in sludge AD from typically 35% to 50% through enhancing structure disintegration of sludge flocs and cell lysis of sludge microorganisms^10,16^. Nonetheless, the sludge pretreatment as a two-edged sword increases concentrations of cell-digestion-derived free ammonia and exacerbates inhibition on sludge AD microbiomes^14,15^, constraining the further improvement of sludge digestion.

In this study, based on an innovative sludge liquefaction and stratification technique, we develop a SPREAD process to boast ultra-high organic loading rates and overcome above-mentioned limitations in sludge AD. We resolve the ammonia inhibition issue through both the prior-to-digestion crude protein harvest from liquefied sludge and the tradeoff of mass concentration and retention time in digester influent, removing the limitation of ammonia inhibition on methanogenic microbiomes and consequent organic loading rates of anaerobic digesters. To test the potential of this novel process, we employ low organic content WAS (0.53 VSS/TSS) as influent and achieve an organic loading rate of 8 kg COD m³ day^-1^ and around 50% N/P recovery, of which performance can be further improved with organic-rich WAS and upgraded digesters. This study provides a game-changing strategy for sludge treatment, resource recovery and sustainable management in WWTPs, of which the concept can be extended to treatment and disposal of other organic-rich solid waste.

## Results

### Liquefaction and stratification of waste activated sludge

Thermal-alkaline sludge pretreatment synergistically combines the individual strengths of thermal- and alkaline-treatments in releasing organic matter from sludge microbial cells (e.g., nucleic acids and proteins) and EPS (e.g., polysaccharides and proteins), respectively^10,17^. Process including TAAD (thermal-alkaline pretreated anaerobic digestion) was developed based on conventional sludge AD to digest the thermal-alkaline pretreated sludge brew with around 30% improvement in organic carbon removal but identical organic loading rates compared to digestion of untreated WAS^10^. In this study, surprisingly, we observed that the thermal-alkaline pretreated sludge brew could be further stratified into three layers after being placed at room temperature in the dark without shaking for 24 hours under an optimized condition (Fig. 1a). Based on their elemental compositions and protein concentrations, the three layers from bottom-to-top were characterized as ash, crude protein and a liquid layer on top (Fig.1). Non-volatile suspended solids (inorganic solids) predominantly accumulated in the ash layer. This resulted in a decrease of VSS/TSS (ratio of volatile suspended solids to total suspended solids) from 0.53±0.01 in the feed WAS to 0.28±0.01 in the ash layer, and released 65.3±2.9% of the organic matter from WAS into the protein and liquid layers (Supplementary Fig.1a). Accordingly, TOC (total organic carbon), TN (total nitrogen) and TP (total phosphorus) in ash accounted for 34.7%, 13.8% and 34.6%, respectively, of their total amounts in the feed WAS (Fig. 1b). In contrast, heavy metals were largely present as precipitates (with a mean value of 72.2% removal efficiency) in the alkaline ash layer (Fig.1c). This largely alleviated their inhibitory effects on subsequent methanogenic digestion and decreased their concentrations in WAS-derived crude proteins and struvite. After ash removal through centrifugation, pH of the liquefied sludge was adjusted from around 11.5 to neutral (pH=7.0) based on the fact that a pH close to the isoelectric points of the proteins would facilitate subsequent protein precipitation^18^. Under the neutral pH condition, protein concentrations in the liquefied sludge decreased from 6.08 g L^-1^ to 3.81 g L^-1^ (Supplementary Fig.1b), with 2.27 g crude proteins retrieved per Litre WAS (30 g L^-1^ TSS or 15.9 g L^-1^ VSS).

**Fig. 1.**
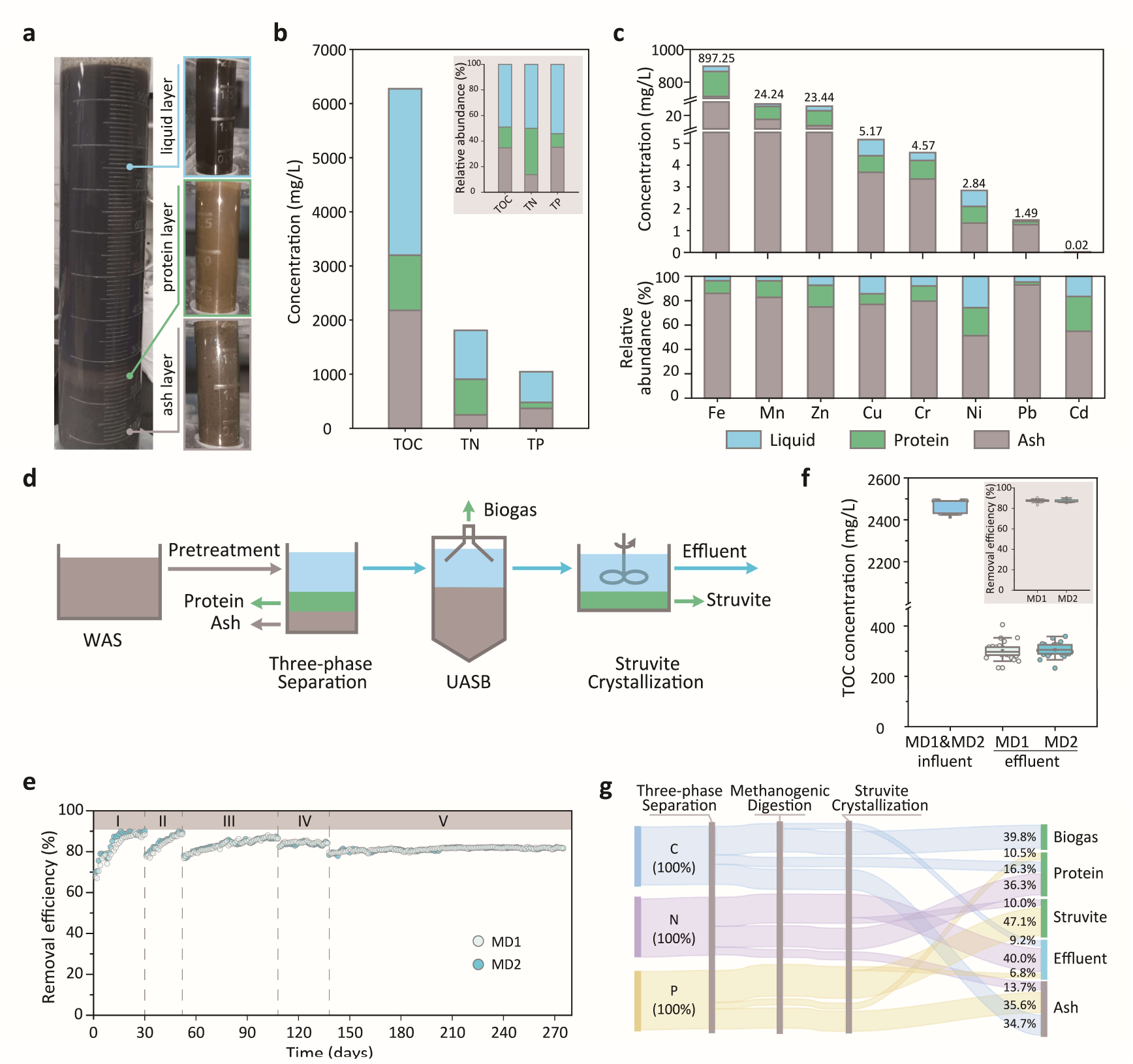
Development of a SPREAD process based on sludge liquefaction and stratification for resource recovery from waste activated sludge. **a,** Stratification of liquefied sludge into liquid, protein and ash layers; **b,** distribution of sludge organic carbon/C, nitrogen/N and phosphorus/P in terms of TOC (total organic carbon), TN (total nitrogen) and TP (total phosphorus), respectively, in the three layers; **c,** the absolute and relative abundance of heavy metals in the three layers; **d,** schematic diagram of the SPREAD process; **e,** TOC removal in duplicate SPREAD UASB (up-flow anaerobic sludge blanket) reactors, MD1 and MD2; f, the influent and effluent TOC concentrations in the duplicate UASB reactors in Stage-V; g, C, N and P resource flows in the SPREAD process.

### Development of SPREAD for exhaustive resource recovery from WAS

For resource recovery from WAS based on the sludge liquefaction and stratification technology, a SPREAD (**<u>s</u>**ludge **<u>p</u>**retreatment and st**<u>r</u>**atification for **<u>e</u>**fficient **<u>a</u>**naerobic **<u>d</u>**igestion) process was developed to retrieve carbon/C, nitrogen/N and phosphorus/P resources as biogas, crude protein and struvite (Fig. 1d). For methanogenic digestion of organic matter in the liquid layer, duplicate up-flow anaerobic sludge blanket (UASB) reactors (i.e., MD1 and MD2) were set-up and operated under identical conditions to convert organic C, N, and P into biogas, ammonium (NH4^+^) and phosphate (PO4^3-^), respectively. Being different from the solid-solid mass transfer and completely mixed digestion microbiome-substrate in AD reactors, the UASB reactor with a three-phase (solid-liquid-gas) separator allows the solid-liquid mass transfer and separation of digestion microbiome from influent substrate, which enable high organic loading rates of the SPREAD process^19–21^. Consequently, organic loading rates in MD1 and MD2 were step-wise increased from 0.6 to 2.4 kg m^-3^ day^-1^ TOC (corresponding to 2-8 kg m^-3^ day^-1^ COD) in five stages through adjustment of influent TOC concentrations, being in line with the increase of effluent concentrations of TN, NH4^+^, TP and PO4^3-^ (Supplementary Fig.2a-d). The mean values of TOC removal efficiency ranged from 88.6±0.9% in Stage-I to 81.2±0.8% in Stage-V (Fig. 1e), with stabilized concentrations of effluent NH4^+^-N and PO4^3-^-P at 556.5±22.7 and 404.0±11.8 mg L^-1^, respectively, in the Stage-V (Supplementary Fig. 2e). Notably, compared to the neutral influent pH, effluent pH fluctuated slightly between 7.4-8.3 (Supplementary Fig. 2f), possibly due to the organic nitrogen-derived NH4^+^ release in the methanogenic digestion process. To test the maximum digestion potential of the influent organic matter in the liquid sludge layer, the organic loading rate was lowered from 2.4 to 0.75 kg m^-3^ day^-1^ TOC (corresponding to 2.5 kg m^-3^ day^-1^ COD) after Stage-V and kept at that rate for 30 days. Accordingly, the effluent TOC in MD1 and MD2 decreased and stabilized at 301.9±32.6 and 306.0±25.6 mg L^-1^, resulting in removal efficiencies of 87.5±1.4% and 87.3±1.0% in MD1 and MD2, respectively (Fig. 1f). These results imply a maximum removal efficiency of 85-90% organic matter in the liquid sludge layer.

To reveal organic matter removal in units of the SPREAD process, excitation-emission matrix (EEM) spectra was employed to monitor organic matter composition profiles in the unite influent and effluent. The EEM results confirmed the removal of proteins and carbohydrates predominantly in crude protein precipitation and methanogenic digestion steps, respectively (Supplementary Fig.3), of the SPREAD process. P/N resources in effluents of the MD1 and MD2 were recovered via struvite crystallization, which resulted in 87.4±1.6% P and 20.0±1.2% N recovery (Supplementary Fig. 4). In the SPREAD process, C/N/P resources in WAS were finally ended up in ash, crude protein, biogas, struvite and final effluent, which lead to an overall of 56.1%, 46.3% and 58.3% recovery of the organic C, N and P resources, respectively, in forms of the crude protein (16.3%C, 36.3%N, and 10.6%P), biogas (39.8%C, 0%N, and 0%P) and struvite (0%C, 10.0%N, and 47.7%P) (Fig. 1g).

### Unique microbial communities and their temporal change in the methanogenic digester of SPREAD

To profile the microbial communities and their changes over time in the duplicate UASB reactors, 16S rRNA gene amplicon sequencing-based analyses were employed to elucidate community succession across the five stages with stepwise increased organic loading rates (Fig. 2 and Supplementary Fig. 5). The outcomes of these analyses were then compared with literature observations on conventional UASB and AD digesters. Principal Coordinate Analysis (PCoA) revealed that the microbial community structure in the SPREAD UASB reactors was distinct from that of conventional anaerobic digesters (Supplementary Fig.5a). With increased organic-loading rates, microbial communities in MD1 and MD2 rapidly shifted in a similar manner and ultimately stabilized in Stage V (Fig. 2a). Surprisingly, microbial source tracking analysis showed that the methanogenic digestion microbiota in the duplicate UASB reactors shared 65.5±10.0% and 28.1±11.7% community similarity with microbiomes of sludge AD and general UASB reactors, respectively (Fig. 2b), suggesting that the influent substrate together with organic loading rate played a key role in driving community assembly in the methanogenic digestion microbiome relative to the reactor configuration and operation mode.

**Fig. 2.**
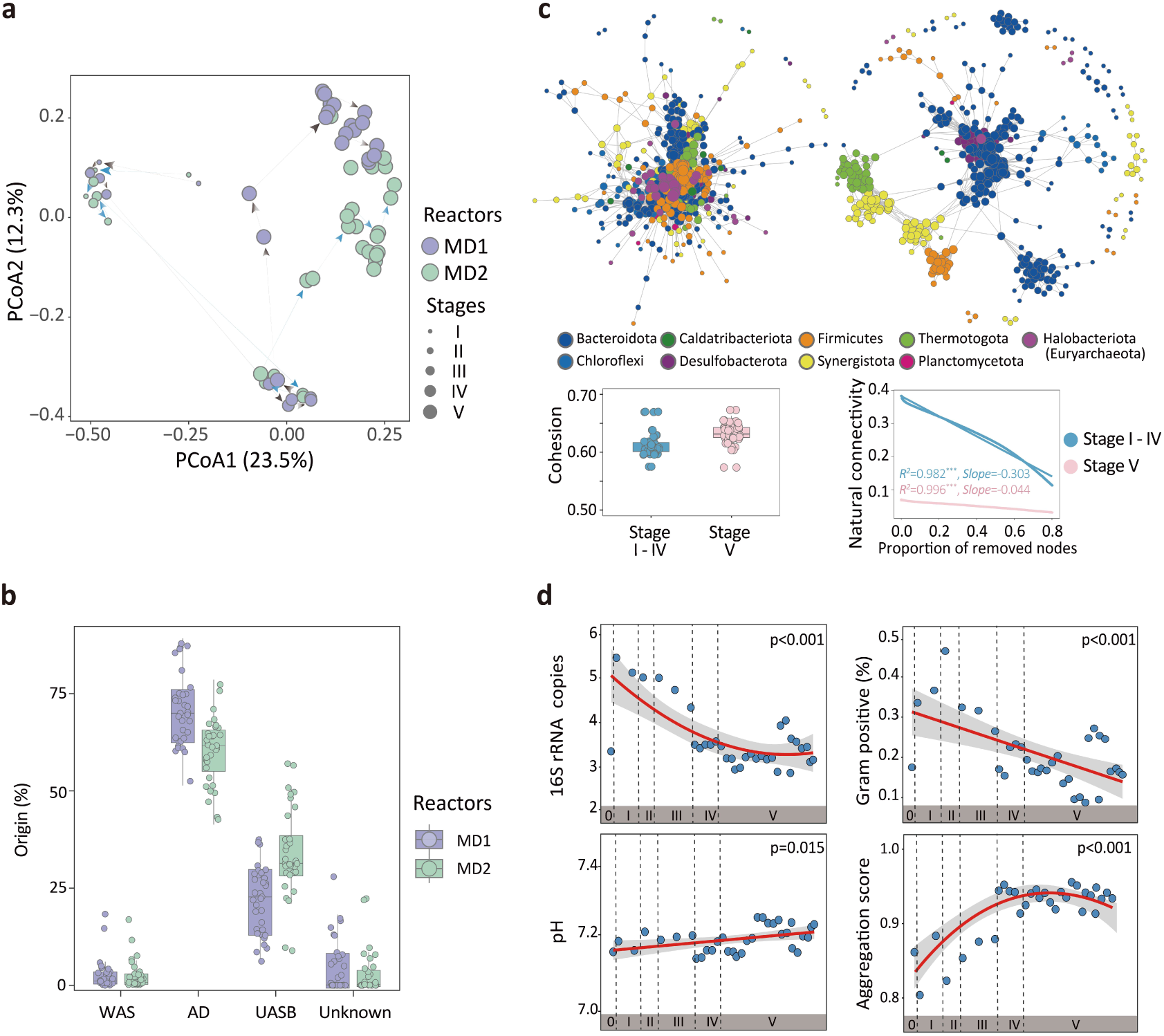
Microbial community succession and source-tracking analyses of the SPREAD UASB microbiomes. **a,** Principal coordinated analysis (PCoA) of the five successive stages of the digestion microbiomes in MD1 and MD2; **b,** microbial source-tracking of Stage-V communities in the duplicate UASB reactors; **c,** comparison of Stages I-IV and Stage-V communities based on Spearman correlation-based microbial co-occurrence network analysis; **d,** predicted 16S rRNA gene copies, pH, Gram-positive bacteria percentage and aggregation score of stages I-V communities in MD1 and MD2. Statistical significance of experimental data (dots) and predicted values (line) are indicated by p-values.

With the increase of organic-loading rates from 2 to 8 kg m⁻³ day⁻¹ COD, the enhanced microbial cohesion and stable natural connectivity indicated improved microbial interactions and community robustness (Fig. 2c), and guaranteed the digestion performance. The improved microbial interactions and community robustness were further corroborated by observing the sludge granulation in stage-V UASB reactors (Supplementary Fig.5b). Further comparison of microbial communities in reactors fed with various feedstocks revealed that increasing the organic-loading rates was associated with improved negative/positive cohesion (Supplementary Fig.5c), reduced sporulation scores (Supplementary Fig.5d) and a decreased number of total genes (Supplementary Fig.5e). These changes contributed to the development of stable, highly specialized and efficient microbial communities in SPREAD. Co-occurrence network analysis indicated that Stage-V communities formed a distinct microbial network compared to the prepositive four stages (Fig. 2c). In Stages I-IV, the network displayed minimal modularization, with closely-clustered key populations of fermenters (Bacteroidota), syntrophs (Desulfobacterota) and methanogens (Halobacteriota) as described previously^22^. In contrast, after 276 days of gradual increase of organic loading rates, communities in Stage-V exhibited a distinct interaction pattern with increased modularity that indicated enhanced stability and resilience^23^. Specifically, only methanogens (*Methanothrix*) and fermenters (*Rikenellaceae*) formed close associations, while other categories of functional microorganisms formed separated clusters (Synergistota, Firmicutes and Thermotogota) (Fig. 2c). To assess trait changes in SPREAD digesters, we developed a comprehensive evaluation and prediction model^24^ by integrating trait data from existing literature with predictions derived from our study (Fig. 2d). Overall, over 60% of the trait-weighted averages exhibited obvious trends during succession, highlighting the predictability of functional development within digestion microbiome. The decrease in 16S rRNA gene copy numbers indicated a transition of the methanogenic digestion microbiomes from rapid-growing and degradation-inefficient cells in floc-like sludge to slow-growing and degradation efficient cells in granular sludge (Supplementary Fig.5b), accompanying improved efficiency in organic matter utilization^25,26^. The predicted pH values aligned with the observed data (*p* = 0.015; Fig. 2d and Supplementary Fig.2f), indicating ammonia release with increased organic loading rates.

### 30-times increase in relative abundance of methanogens rationalize the ultra-high organic loading rates in SPREAD

To gain insight into the mechanism behind the ultra-high organic loading rates of SPREAD relative to conventional sludge AD, microbial community compositions in SPREAD UASB reactors were compared to these of other UASB and conventional AD fed with varied-energy-extent substrates, i.e., high-energy carbohydrates (glucose, maltose and sugarcane), low-energy VFAs and terephthalic acid, and medium-energy organic mixtures (WAS, straw and food waste) (Fig. 3 and Supplementary Table 1). Surprisingly, in contrast to the 1.1% in relative abundance of total methanogens in AD (TAAD) fed with similar pretreated WAS without sludge liquefaction and stratification, relative abundance of total methanogens achieved 33.5% in the SPREAD UASB (Fig. 3a and b). The 30 times increase in relative abundance of the methanogens can rationalize the organic loading rates of 8 kg m^-3^ day^-1^ COD in the SPREAD compared to 0.5 kg m^-3^ day^-1^ COD in the TAAD. Interestingly, further comparisons of the slow-growing methanogens showed their low relative abundance (2.6±1.4%) in all AD microbiota, in contrast to their increasing relative abundance in UASB microbiota along with the substrate-energy increase, i.e., 73.2±7.5%, 53.4±11.7% and 6.5±5.4% in UASB fed with VFAs, organic mixtures and carbohydrates as influent substrates (Fig. 3b and Supplementary Table 1). This observation suggested that the slow-growing methanogens could be hardly retained in AD microbiome due to its completely mixed with influent substrate, frequent biomass discharge and consequent identical hydraulic retention time (HRT) and sludge retention time (SRT). The significantly higher 16S rRNA gene copy numbers of the AD microbiomes compared to the UASB microbiomes further support this, since microorganisms with high 16S rRNA gene copy numbers generally have short cell generation time and rapid growth (Fig.3c).

**Fig. 3.**
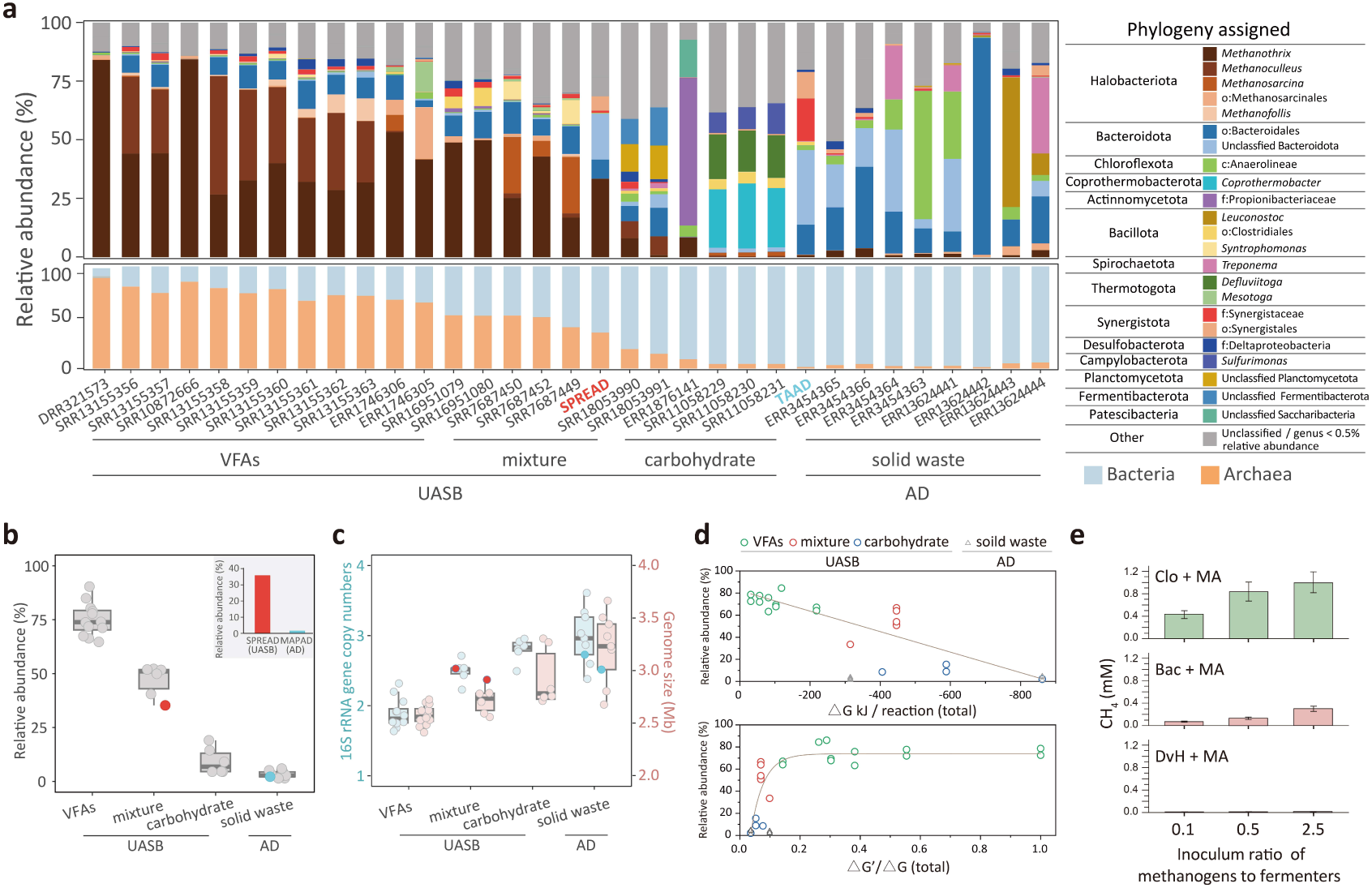
Correlation of relative abundance of methanogens with the energy content of various substrates in methanogenic digestion microbiomes. **a,** Microbial community composition and relative abundance of methanogens in microbiomes of UASB (HRT and SRT are different) and AD (HRT and SRT are identical) reactors fed with low-energy volatile fatty acids (VFAs) or terephthalic acid, high-energy carbohydrates, or medium-energy organic mixture of proteins and lipids or solid wastes; **b,** the relative abundance of methanogens in the communities shown in Fig.3a; **c,** predicted 16S rRNA gene copy numbers and genome size of the communities shown in Fig.3a; the communities of SPREAD UASB and TAAD AD reactors fed with similar sludge were marked in red and blue circles; **d,** correlation between the total Gibbs free energy (ΔG) of methanogenic digestion of varied-energy-extent substrates and the relative abundance of methanogens, and between the ratio of methanogenesis-derived energy to total Gibbs free energy (ΔG m /ΔG) and the relative abundance of methanogens (see Supplementary Table 1 for detailed equations); **e,** impact of gradient bioaugmentaion of methanogens (0.1, 0.5 and 2.5 inoculum ratio of the methanogen to fermenter) on methane generation in different co-cultures fed with varied-energy-extent substrates: Clo+MA, a co-culture of *Clostridium*/Clo and *Methanosarcina*/MA fed with high-energy glucose; Bac+MA, a co-culture of Bacteroides/Bac and MA fed with high-energy glucose; DvH+MA, a co-culture of *Desulfovibrio*/DVH and MA fed with low-energy In the UASB where SRT and HRT are separated^19,20^, very interestingly, the relative abundance of methanogens was negatively correlated with the Gibbs free energy (*Δ*G) derived from methanogenic digestion reactions of influent substrates (R² = 0.771, *p* < 0.001), in contrast to no correlation between *Δ*G and methanogens’ relative abundance in AD microbiomes (Fig. 3d and Supplementary Table 1). Moreover, correlation between the relative abundance of methanogens and the proportion of methanogenesis-derived *Δ*G fitted the Boltzmann equation (R² = 0.738, *p* < 0.001), and the relative abundance of methanogens peaked at 65% when the methanogenesis-derived *Δ*G accounted for around 15% (e.g., methanogenic digestion of maltose) of the total methanogenic-digestion-derived *Δ*G (Fig. 3e and Supplementary Table 1). Therefore, the separation of SRT and HRT in UASB enabled the correlation between the relative abundance of methanogens and the substrate energy derived from anaerobic digestion.

These results further indicated that the low relative abundance of methanogens was the predominant efficiency-limiting factor in digestion of high-energy substrates, and augmentation of methanogens could enhance the methanogenic digestion efficiency. To further test it, three sets of experiments were setup with three fermenting/acetogenic bacteria (*Clostridium*/Clo, *Bacteroides*/Bac and *Desulfovibrio*/DvH) and a methanogen (*Methanosarcina*/MA; Fig. 3f): (1) Clo-MA, coculture of *Clostridium* and *Methanosarcina* with glucose as a high-energy substrate; (2) Bac-MA, coculture of *Bacteroides* and *Methanosarcina* with glucose as a high-energy substrate; (3) DvH-MA, coculture of *Desulfovibrio* and *Methanosarcina* with lactate as a low-energy substrate. In contrast to no augmentation effect of adding extra amounts of methanogens in the DvH-MA coculture fed with a low-energy substrate, methanogenic activities were linearly and positively correlated with the augmented methanogens fed with a high-energy substrate (Fig. 3f). These results corroborated that abundance of methanogens could be a key factor and bottleneck to determine the efficiency in methanogenic digestion of high-energy substrates, providing a strategy for bioaugmentation of UASB microbiomes.

### Meta-omics reconstruction of metabolic networks in methanogenic digestion microbiome of SPREAD

To comprehensively assess the metabolic network in the SPREAD UASB microbiome, an anaerobic digestion database (ADDB) was developed based on both top-down and bottom-up strategies to facilitate meta-omics analyses (Fig. 4 and Supplementary Fig.6). For the top-down data curation, pathways associated with digestion metabolism were identified based on the literature and related databases (KEGG and MetaCyc)^27,28^. To validate and complement the data derived from the top-down strategy, bottom-up data curation was manually performed, beginning with metabolites and intermediates from the MetaCyc and KEGG databases (Supplementary Fig. 6). Then, directed metabolic networks were constructed with representative metabolites as nodes and their transformation processes (enzymatic reactions) as edges with the depth-first search method^29^. The integration of top-down and bottom-up strategies ensured the accuracy and coverage of ADDB for meta-omics analyses. Overall, a total of 4,588 genes and gene-encoded enzymes, 1,216 reactions, and 1,268 metabolites were included, and encompassed 4 modules (level-1) and 24 sub-modules (level-2) related to depolymerization/hydrolysis, fermentation, acetogenesis, methanogenesis, and other digestion-related metabolic processes, e.g. WLP-based acidogenesis and carbon-chain elongation (Fig. 4a).

**Fig. 4.**
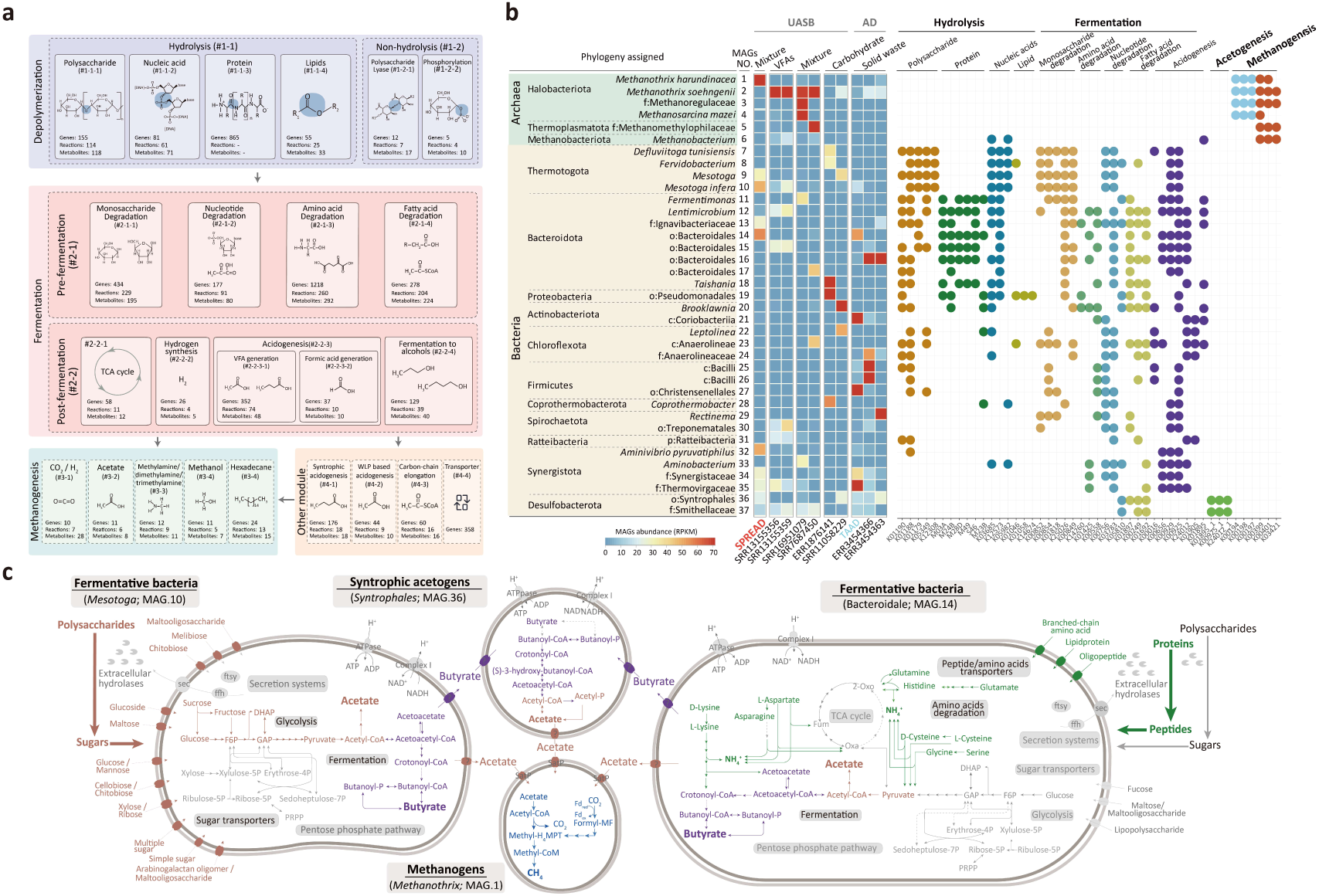
Meta-omics reconstruction of metabolic networks in methanogenic digestion microbiome of SPREAD. **a,** Modularized content of the anaerobic digestion database (ADDB); **b,** MAGs-based taxonomic and functional profile comparison of digestion microbiomes fed with varied-energy-extent substrates in UASB and AD; **c,** schematic representation of intra- and inter-cellular metabolic networks of the major functional populations in SPREAD UASB microbiome, i.e. fermenters (*Mesotoga* and Bacteroidales), syntrophic acetogens (Syntrophales) and methanogens (*Methanothrix*). Colors highlight the metabolic pathways confirmed with metatranscriptome and metaproteome data (see Supplementary Figure 9 and Table 5 for detailed metatranscriptomic and metaproteomic profiles).

In the metagenomic analysis of SPREAD UASB microbiome, 31 bacterial and 6 archaeal MAGs (metagenome-assembled genomes) were retrieved from 35 Gb of total raw sequencing reads (Supplementary Table 2, 3, and 4), which spanned 22 phyla and involved all of the major methanogenic digestion processes (Supplementary Fig.7). A comparison of the major methanogenic digestion players in the SPREAD UASB microbiome and other digestion microbiomes showed that their community compositions were predominantly determined by the influent substrate (VFAs, organic mixture and carbohydrates as low-, medium- and high-energy substrates, respectively) and reactor configuration (UASB *vs.* AD) (Fig.4b and Supplementary Fig.8). In contrast to their different fermenting bacteria, UASB microbiomes fed with low- and medium-energy substrates were all dominated by identical genus of methanogens (*Methanothrix*; Fig.4b). Specifically, fermenting bacteria in the SPREAD UASB microbiome were mainly composed of polysaccharide- and nucleic-acid-fermenting *Mesotoga* (MAGs #9 and #10) and protein/nucleic-acid/polysaccharide-fermenting Bacteroidales (MAGs #13 and #14) (Fig.4b and Supplementary Fig.8a). Further metatranscriptomic and metaproteomic analyses of the SPREAD UASB microbiome further confirmed the central roles of these populations in methanogenic digestion of the sludge-derived organic matter (Supplementary Fig.8b and c). Interestingly, though populations (MAGs #13 and #14) of the Bacteroidales order were showed to have the potential in fermentation of proteins, nucleic acids and polysaccharides based on the metagenomic analysis (Fig.4b and Supplementary Fig.8a), their metatranscriptomic and metaproteomic data indicated that they were specifically and highly active in fermentation of the WAS-derived proteins in SPREAD (Fig.4c and Supplementary Fig. 8b and c). Therefore, populations of the Bacteroidales and *Mesotoga* were complementary to each other in fermentation of proteins and nucleic-acids/polysaccharides, respectively, which further formed metabolic networks with acetogenic *Syntrophales* and methanogenic *Methanothrix* for extensive digestion of WAS-derived organic matter in SPREAD.

### Environmental impacts and economic benefits of SPREAD

A Life Cycle Assessment (LCA) was performed to evaluate the environmental impact and economic benefit of the SPREAD process in comparison with two commonly used anaerobic digestion processes: conventional anaerobic digestion (CAD) and anaerobic digestion with thermal-alkaline pretreatment (TAAD) (Fig. 5a; Supplementary Table 7). SPREAD outperformed the other two processes across all 14 impact categories (Fig. 5b). Among those, SPREAD achieved the most significant benefits in human toxicity potential and freshwater eutrophication potential, with reductions of 44.9% and 60.8% compared to CAD, and 71.9% and 78.8% relative to TAAD, respectively. The improves are primarily attributed to the efficient N/P recovery, which alleviated subsequent WWTPs-effluent-derived surface water eutrophication. Pertaining to the carbon footprint, SPREAD was demonstrated to have obvious advantages over CAD and TAAD, i.e., 15.0, 28.5, and 33.9 kg CO2 eq per ton of sludge (97% moisture content) of the SPREAD, CAD and TAAD, respectively (Fig. 5c).

**Fig. 5.**
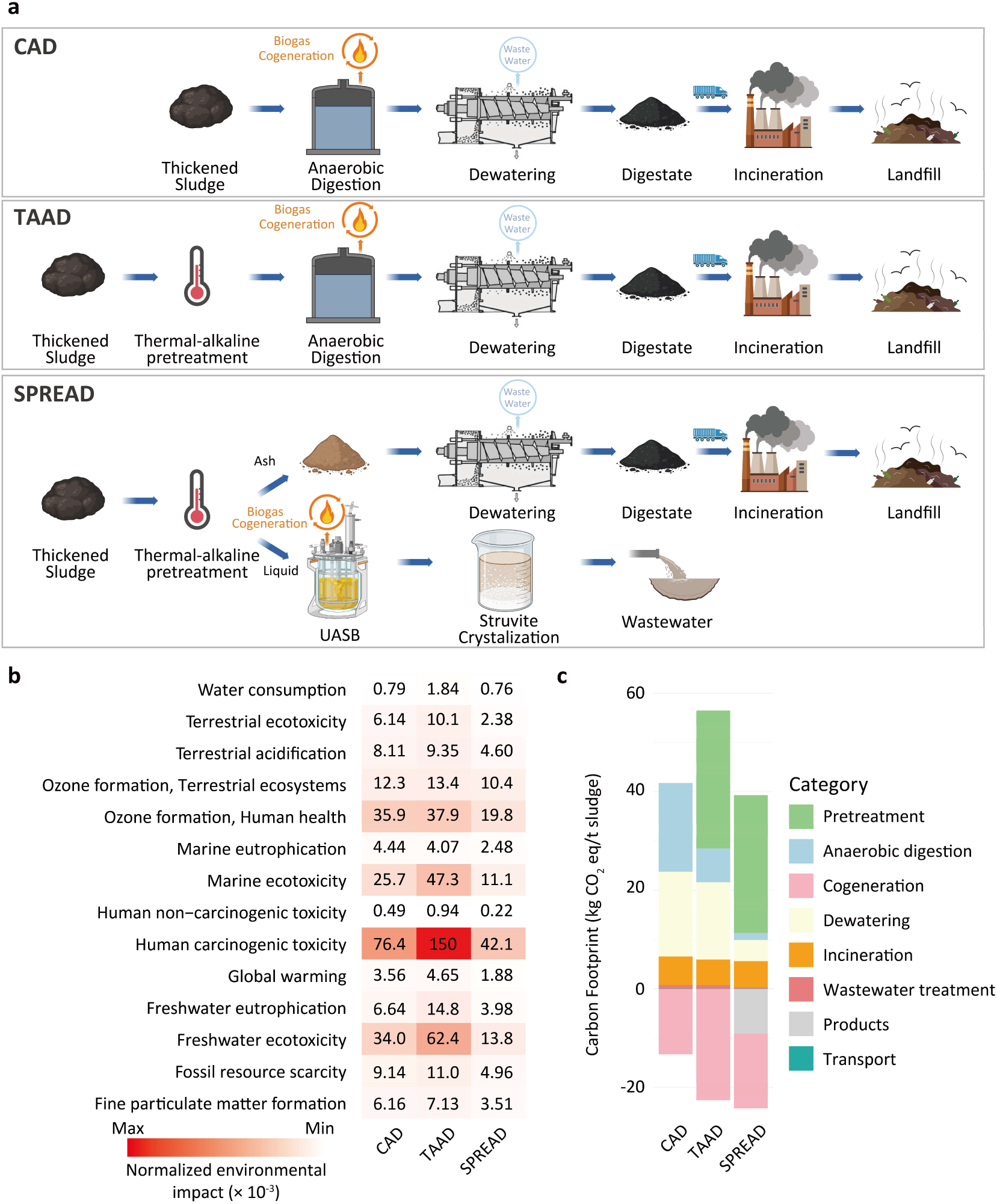
Life cycle environmental impact and economic assessment of SPREAD. **a,** System boundary of conventional mesophilic anaerobic digestion (CAD), anaerobic digestion combined with thermal-alkaline pretreatment (TAAD), and SPREAD; **b,** Hotspots of 14 environmental impact categories. The color scale ranks the magnitude of the impacts (with red denoting a high impact); **c,** Carbon footprint per ton sludge (97%, moisture content) of CAD, TAAD, and SPREAD

## Discussion

In this study, a novel process entitled SPREAD was developed for exhaustive energy and N/P resource recovery from WAS based on an innovative technique of sludge liquefaction and stratification into three (ash-protein-liquid) layers. The process achieved approximately 10 times higher organic loading rates relative to conventional sludge AD process (Table 1), which could reduce the land-footprint of current sludge digesters and WWTPs by 10 times and 30%, respectively^31,32^. The extraordinary performance of the SPREAD was mainly due to the sludge liquefaction and stratification for overcoming limitations of the conventional sludge AD: (1) sludge liquefaction led to dissolution of organic matter from the solid-form WAS, improving mass transfer between substrates and microorganisms, i.e. solid-solid mass transfer in conventional AD *vs*. liquid-solid mass transfer in SPREAD. Importantly, it also allows the separation of HRT (hydrolytic retention time) from SRT (sludge retention time) in digesters, which is one of the defining characteristics of the successful UASB technology. In addition, mass transfer between substrates and microorganisms is generally the key rate- and efficiency-limiting parameter in bioreactors^11,12,33^, e.g., the organic loading rate could be increased from 10 kg m^-3^ day^-1^ COD in UASB reactors (microorganisms in sludge bed) to 50 kg m^-3^ day^-1^ COD in IC (internal circulation reactor; microorganisms in fluidized sludge bed) reactors by fluidizing the sludge bed and consequently improving the mass transfer coefficient^34^. In conventional sludge AD, the solid-form sludge is digested and discharged with active methanogenic digestion biomass, which results in the coupled and identical long HRT and SRT (20-30 days) and the biomass loss of slow-growing methanogens^10,35^. In contrast, SPREAD with liquefied WAS as influent enables the separation of HRT (12 hours) from SRT (6 months), allowing 30 times higher in biomass abundance of slow-growing methanogens compared to conventional sludge AD. (2) ammonia inhibition associated with conventional sludge AD was alleviated in the SPREAD process by harvesting around 40% WAS-derived crude proteins from the ash-protein-liquid layers and by balancing the tradeoff between influent COD concentrations and HRT for adjustment of the organic loading rate. With a specific organic loading rate, the shorter HRT enables the lower concentration of influent organic matter in the SPREAD process, which prevents organic-derived and concentration-dependent ammonia inhibition^13,15^. (3) the organic loading rate for sludge digestion in SPREAD was increased for over than 10 times based on the synergistic effects of above-mentioned properties, which could be further improved by employing IC and other upgraded reactors^34,36^. Moreover, the SPREAD process addresses a fundamental challenge in biogas separation and collection of conventional AD. Carbon footprint analyses of WWTPs have indicated that biogas leakage from anaerobic digesters and subsequent digested-sludge processing steps contributes approximately 37.5% of the total carbon emissions^37^. SPREAD prevents the biogas leakage through the more effective gas-liquid separation, resulting in a reduction of carbon emissions by approximately 60% compared to conventional AD. This enhancement offers substantial improvements across all LCA metrics of the SPREAD. Consequently, the SPREAD process represents a potentially game-changing solution for WAS treatment, addressing the intertwined challenges of resource recovery, environmental protection, and cost-efficiency in WWTPs.

**Table 1.** Comparison of SPREAD and conventional sludge AD.

| Technical parameters | Conventional AD | SPREAD |
| --- | --- | --- |
| Feedstock | WAS | WAS |
| Hydrolytic retention time (HRT) | 20-30 days | 8-12 hours |
| Sludge retention time (SRT) | 20-30 days | ≥6 months |
| Organic loading rate ($\text{kg m}^{-3} \text{ day}^{-1}$ COD) | 0.5-1 | 8 |
| Organic carbon recovery | 35% | 56% |
| Nitrogen recovery | <10% | 46.3% |
| Phosphorus recovery rate | <10% | 58.3% |
| Ammonia inhibition | Yes | No |

In well-controlled biosystems such as UASB and AD reactors, the microbial community diversity, stability and their succession are fairly predictable, and less diverse communities are generally more stable^38,39^. In contrast to the complex communities in conventional sludge AD, methanogenic communities in the SPREAD UASB were dominated by *Mesotoga*, Bacteroidales, *Syntrophales* and *Methanothrix*.

These four groups accounted for >65% of the total microbial communities, which supported the stable UASB microbiome with high organic loading rate. According to the “streamlining theory”, microbial communities subjected to environmental stress (or selective pressure, e.g. high organic loading rate) tend to have reduced functional redundancy^40^, which could be helpful to improve the community efficiency and stability^39,41^. Therefore, the streamlining theory well rationalized the SPREAD UASB microbiome with enrichment of major functional microorganisms and enhancement of community stability and consequent organic loading rate. The exceptionally high relative abundance of the slow-growing acetoclastic methanogen *Methanothrix* was 30 times higher in the SPREAD UASB compared to the conventional sludge AD, which can be rationalized by the above-mentioned ecological principle and following two reasons: (1) decoupling of HRT (substrate retention time) and SRT (biomass retention time) ensured retaining high relative abundance of the slow-growing *Methanothrix*; (2) short-HRT-enabled low concentrations of influent substrates prevented ammonium accumulation and consequent ammonia inhibition on *Methanothrix*.

Our observation of the negative correlation between the Gibbs free energy (*Δ*G) and the relative abundance of methanogens may offer a new tool for the identification of rate- or efficiency-limiting steps for stimulation and augmentation of methanogenic digestion microbiomes: (1) for methanogenic digestion reactions with low *Δ*G but high relative abundance of methanogens, the acidogenic fermentation could be the rate-limiting step; and (2) for the methanogenic digestion reactions with high *Δ*G but low relative abundance of methanogens, the biomass of methanogens could be a rate- and efficiency-limiting factor. In previous studies, conclusions on the rate- or efficiency-limiting steps have been controversial, which may negatively affect the setup, maintenance and optimization of methanogenic digestion microbiomes^42–44^. The key reason underlying the controversial debate was the failure to appreciate the negative correlation between the *Δ*G and the relative abundance of methanogens. Therefore, our observation opens a new avenue for assessing the bottleneck step in methanogenic digestion, which could advance our understanding and operation of the methanogenic digestion biosystems in general, beyond mere sludge digestion systems.

## Methods

### Sludge liquefaction and stratification

Waste active sludge (WAS) was collected weekly (from July 2020 to June 2021) from a secondary clarifier of a local wastewater treatment plant (WWTP) in Guangzhou, China. Prior to subsequent pretreatment, the sludge was concentrated to a final total suspended solids (TSS) concentration of 30 g L^-1^. The concentrated sludge was then subjected to thermal-alkaline pretreatment by addition of 0.15 M NaOH, followed by incubation at 170°C for 70 min in sealed bottles as described^10^. After pretreatment, three distinct layers (i.e. ash, crude protein and liquid layers) were observed after setting under room temperature (Fig. 1a). These layers were subsequently separated using the following procedures: (i) the ash layer was collected by centrifuging the sludge mixture at 3000 r min^-1^ for 3 min to separate it from crude protein and liquid layers; (ii) the crude protein and liquid sludge layers were neutralized to a pH of 7.0 by adding HCl, after which the crude protein layer was collected by centrifugation at 4000 r min^-1^ for 5 mins; (iii) the liquid layer of liquidized sludge was utilized as influent substrate for methanogenic digestion in the SPREAD UASB reactor.

### Development of a SPREAD process for exhaustive resource recovery from WAS

Based on the observation of sludge liquefaction and stratification, a SPREAD (**<u>s</u>**ludge **<u>p</u>**retreatment and st**<u>r</u>**atification for **<u>e</u>**fficient **<u>a</u>**naerobic **<u>d</u>**igestion) process was developed to recover carbon(C), nitrogen(N), and phosphorus(P) resources from WAS in the forms of biogas, crude protein, and struvite (Fig. 1b). In the anaerobic digestion section, duplicate up-flow anaerobic sludge blanket (UASB) reactors (i.e. MD1 and MD2) with a working volume of 2 L were set-up for the treatment of the liquid layer of liquidized sludge. Methanogenic digestion sludge as reactor inoculum was collected from a sludge digester^45^. The hydraulic retention time (HRT) was maintained at 24 h, and the reactor operation temperature was kept constant at 35 ± 0.5°C using a water bath. The organic loading rates were increased stepwise from 0.6 to 2.4 kg m^-3^ day^-1^ TOC (corresponding to 2-8 kg m^-3^ day^-1^ COD), i.e., 0.6, 1.2, 1.8, 2.1 and 2.4 kg m^-3^ day^-1^ TOC in Stage-I, II, III, IV and V, respectively. Nutrient recovery from digestion effluent (pH=8) was achieved through magnesium chloride addition and subsequent struvite crystallization, with an optimized Mg/P molar ratio of 1.2:1^46^.

### Analytical methods

The elemental compositions of suspended solids were analyzed with a CHON elemental analyzer (Vario EL, Elementar; GmbH, Germany)^45^. Heavy metal concentrations were tested using an inductively coupled plasma optical emission spectrometry (ICP-OES, 5300DV, Perkin Elmer, USA) as described^47^. Excitation-emission matrix (EEM) spectra of organic matter in the liquidized sludge, and digester influent and effluent were obtained using a LS-55 spectrofluorometer (Perkin-Elmer, USA) following a described method^48^. Briefly, excitation wavelength was set in the range of 200-400 nm with 10 nm intervals, and the emission wavelength was measured between 200-550 nm with 0.5 nm intervals. Ultrapure water was used as a reference to eliminate background noise. Protein concentration was measured according to the Lowry method^10,49^. The TOC, COD, TN, NH4^+^-N, TP, PO4^3-^-P, TSS and volatile SS (VSS) were analyzed following standard methods^50^.

### DNA extraction, amplicon sequencing and data processing

Total genomic DNA was extracted using a TINAamp soil DNA kit (Tiangen, Beijing, China), according to the manufacturer’s instructions. The hypervariable V4-V5 regions of the 16S rRNA genes were amplified with a 515F/909R primer set as described^51^. The purified PCR products were pooled and sequenced on an Illumina Novaseq platform (PE250; Illumina; San Diego, CA, USA) provided by MAGIGENE (Shenzen, China). The demultiplexed paired-end (2×250 bp) reads were further processed using the DADA2 method (version 1.6)^52^ in R software (version 4.2). Taxonomy was assigned by using RDP naïve Bayesian classifier^53^ with the SILVA v138 database^54^. To minimize the bias of sequencing depth, ASVs table was rarefied to 11,800 sequences per sample. Principal coordinates analysis (PCoA) based on Bray-Curtis dissimilarity matrix was performed at the ASV level to reveal the community difference.

### Source-tracking analysis, microbial network and trait prediction of digestion microbiota

Microbial source tracking analysis was conducted using the FEAST method^55^ to estimate the origin of targeting microbial communities. A highly efficient expectation-maximization-based method was applied to assess the contribution of source communities to a sink community^56^. A total of 525 microbiota data from activated sludge, sludge anaerobic digestion, and upflow anaerobic sludge blanket (UASB) were used as source communities (Supplementary Table 6), whereas SPREAD samples were set as sink communities. These 525 microbiota datasets were retrieved from NCBI’s Sequence Read Archive (SRA) database and processed using DADA2 (v3.16)^52^ to obtain the ASV table. The relationship between source and sink was observed using the FEAST package in R software^55^.

For co-occurrence network analysis, only the ASVs presented in at least 20% of all samples were included to reduce dataset complexity. Networks were inferred by the SparCC method^57^, with 1000 permutations tested to determine the correlations between taxa. Only robust (|r|>0.5) and statistically significant (p < 0.01) correlations were considered. Network visualizations were generated within the R environment using igraph packages. Cohesion and network robustness were performed to evaluate microbial community stability. Cohesion quantifying the community connectivity based on the association between taxa and their abundance was calculated by following the protocol as described^58^. Briefly, correlations between all taxa within a community were calculated using Pearson correlation, and a null model was employed to verify the strength of these correlations. Expected correlations derived from the null model were then subtracted from the observed correlations to obtain positive and negative connectedness values. These connectedness values were weighted by the relative abundance of taxa and summed to yield positive and negative cohesion metric. Communities with lower positive cohesion values tended to be more stable^23,58^. Network robustness was assessed as a measure of network stability by examining how natural connectivity changes when nodes or edges were removed in descending order of their betweenness or weight^59^. The natural connectivity of the network was estimated by systematically “attacking” nodes or edges^60,61^. The fastNC software^59^ was used to test the network robustness with the default parameter.

The traits of digestion microbiota were predicted according to the protocol reported by Guittar et al. (2019)^24^. Briefly, a phylogeny was constructed that included taxa from digestion samples along with the long-term persistence (LTP) taxa^62^ with formally described type specimens and Latin binomials. Trait data curated from literature and online repositories were mapped onto the phylogeny tips using the corresponding Latin binomials. Unknown trait values were inferred through hidden state prediction. The aggregation score ranged from 0 to 1, meaning never observed aggregation to consistently observed aggregation. Similarly, the Gram-positive score ranged from 0 (Gram-negative) to 1 (Gram-positive), as well as the sporulation score from 0 (never sporulates) to 1 (sporulates easily).

### Development of the anaerobic digestion database (ADDB)

To facilitate metabolic profiling of anaerobic digestion processes based on meta-omics data, the Anaerobic Digestion Database (ADDB) was developed using a comprehensive approach that combines top-down and bottom-up strategies (Supplementary Fig.6). The ADDB intergrades a wide array of functional genes, metabolites, and reactions involved in the methanogenic digestion, systematically organized into hierarchical modules and sub-modules that reflect steps of methanogenic digestion, including depolymerization/hydrolysis, fermentation, acetogenesis, methanogenesis and other digestion-related metabolic processes (e.g., WLP based acidogenesis and carbon-chain elongation) (Fig. 4a). In the top-down curation process, anaerobic digestion-related metabolic pathways were initially identified from literatures and databases, e.g. KEGG and MetaCyc^27,28^. These pathways were then broken down into specific processes, including depolymerization, fermentation, methanogenesis and others. Key reactions, genes, and metabolites associated with each process were extracted and curated into the ADDB core database. To complement the top-down strategy, a bottom-up approach was employed to ensure comprehensive coverage of metabolic pathways. This approach began with identifying metabolites and then extending the analysis to genes and enzymes involved in these metabolic transformations. By constructing a directed metabolic network where metabolites served as nodes and enzymatic reactions as edges, the entire metabolic landscape of anaerobic digestion was mapped out. The depth-first search method^29^ was applied to explore these networks, starting from initial substrates and tracing through to final products, ensuring that all relevant reactions and pathways were included. Manual curation was also employed to refine the automatically generated networks, ensuring accuracy and relevance to anaerobic digestion. Gene and protein sequences corresponding to the curated pathways were retrieved from databases such as KEGG, UniProt^63^ and MEROPS^64^, ensuring that the ADDB includes detailed sequence information for further functional analyses. Special modules for transporters and other specific functional aspects of anaerobic digestion were also curated to support in-depth metabolism studies.

### Metagenomic, metatranscriptomic and metaproteomic analyses

Sludge samples for metagenomic analysis were collected from MD1 on day 265 (Stage V). Genomic DNA extraction was performed as described above. The quality and quantity of the extracted gDNA were assessed using gel electrophoresis and a Quantus fluorometer (Promega, Madison, WI, USA). Library preparation and metagenomic sequencing on a HiSeq platform (Illumina) were conducted by MAGIGENE (Shenzhen, China). Raw metagenomic sequencing data were filtered to remove low-quality bases/reads using trim_galore (v0.6.10) with default parameters. Metagenomic reads and reference genome were taxonomically profiled using mOTU v3.0.382 and counts were distributed to GTDB species using the GTDB_v214 mapping file, being available as parts of mOTUs database^65^. Each metagenome was assembled using metaSPAdes (v3.10.0)^66^ with k-mer values of 33, 77, and 99. Genome reconstruction of the SPREAD UASB microbiome was performed using the function modules of metaWRAP (v1.2)^67^. Specifically, contigs longer than 1000 bp were used for genome binning. Metagenome-assembled genomes (MAGs) were constructed from contigs in each sample using three different binning algorithms: ’--metabat2 --maxbin2 --concoct’ within the metaWRAP software. Bin refinement was performed using the bin_refinement module of metaWRAP. To achieve optimal genome quality, metagenomic sequencing reads were mapped to each bin and reassembled using metaSPAdes via the reassemble_bins module of metaWRAP. The completeness and contamination of each bin were evaluated using CheckM^68^. dRep v3.4.3^69^ was used to dereplicate high- and medium-quality MAGs (completeness ≥50% and contamination ≤10%) at 95% ANI. MAGs were taxonomically assigned using GTDB-tk v2.1.0^70^. Maximum-likelihood phylogeny of MAGs was inferred using IQ-TREE v2.2.0.3 from a concatenation of 40 marker genes searched MAGs by fetchMGs v1.1^71^. The generated tree was visualized using iTOL v6 (https://itol.embl.de). Functional annotations were conducted based on the ADDB database using DIAMOND^72^ with an E-value threshold of 1e-5. Potential metabolic modules in the metagenome were classified according to ADDB. To reconstruct metabolic pathways, the predicted genes of MAGs were further annotated using the KEGG Automatic Annotation Server (KAAS)^73^ and dbCAN3^74^. The abundance of genes and MAGs was estimated using CoverM (v0.6.0, https://github.com/wwood/CoverM/), which was normalized as reads per kilobase per million (RPKM).

Both metatranscriptomic and metaproteomic analyses were conducted to confirm the transcription and translation of functional genes in SPREAD UASB reactors. Briefly, samples for total RNA and protein extraction were collected on Day 265 by centrifugation (5 min, 10,000×g, 4°C) from triplicate samples. RNA samples were then mixed with TRIzol™ reagent (Invitrogen, Waltham, MA, USA). Total RNA was extracted using the RNeasy Mini kit (Qiagen), and fragmented DNA was subsequently removed with DNase I treatment. Highly transcribed ribosomal RNAs (rRNAs) were depleted using the RiBoCop rRNA Depletion Kit (Lexogen, Vienna, Austria). RNA-Seq libraries were constructed from the rRNA-depleted RNA samples using the NEBNext® Ultra™ Directional RNA Library Prep Kit (New England Biolabs, Ipswich, MA, USA) and sequenced on an Illumina Novaseq6000 platform. RNA-Seq library construction and sequencing services were provided by MAGIGENE (Shenzhen, China). RNA-Seq raw data in FASTQ format were processed using Trimmomatic (v0.35)^75^ to obtain clean reads. The clean reads were first mapped to previously established MAGs to remove rRNA sequences using Bowtie2 (v2.33)^76^. Gene read counts and functional information were obtained using HTSeq-count (v0.9.1) based on the mapping results^77^. RPKM (reads per kilobase of transcript per million mapped reads) values were calculated to compare the expression levels of different genes^78^. For metaproteomic analysis, samples were resuspended in 10 mM phosphate buffer solution (pH 7.4) and centrifuged again prior to storage at -80°C. Further protein extraction, digestion and quantification services were provided by the Beijing Genomics Institute (BGI, Shenzhen, China) as described^79^.

### Life cycle environmental impact assessment of SPREAD

Life cycle assessment (LCA) was performed to compare environmental impacts of SPREAD and two other commonly used sludge digestion processes, i.e. conventional anaerobic digestion (CAD) and anaerobic digestion with thermal-alkaline pre-treatment (TAAD). The system boundaries for the LCA analyses were defined to include the following units: pretreatment, anaerobic digestion, biogas cogeneration, dewatering, and the management of the digestate for incineration and landfill^9,30^ (Supplementary Fig. 9). The functional unit was treating 1 ton of waste sludge with a 97% moisture content. Input and output data were sourced from both literature and experimental results (Supplementary Table 7)^30,80^. Background data were obtained from the Ecoinvent v3.5 database. The LCA was conducted in SimaPro (v9.5.0.0) using ReCiPe midpoint method (v1.13) ^81^.

## Data availability

The authors declare that the data supporting the findings of this study are available within the article and its supplementary information. Illumina sequencing data for this study has been deposited to the European Nucleotide Archive (ENA) database with accession numbers PRJEB79622.

## Acknowledgements

This study was financially supported by the National Natural Science Foundation of China (42161160306 and 42177001) and the Natural Science Foundation of Guangdong Province (2018B030314012).

