## Supplemental information for "Sludge liquefaction and stratification enable ultra-high digestion rate and resource recovery for waste activated sludge treatment"

### These authors contributed equally

*Corresponding authors:

Shanquan Wang

**This file includes:**

**Supplementary Figures 1-9**

**Supplementary Fig. 1 |** Characterization of the liquefied sludge: **a,** VSS/TSS and TSS of waste activated sludge (WAS) and ash layer of the liquefied sludge; **b,** Protein concentrations in the crude-protein and liquid layers of the liquefied sludge.

**Supplementary Fig. 2 |** Performance of the SPREAD UASB reactors MD1 and MD2: **a,** effluent TN concentrations in the five stages; **b,** effluent $NH_4^+$-N concentrations in the five stages; **c,** effluent TP concentrations in the five stages; **d,** effluent $PO_4^{3-}$-P concentrations in the five stages; **e,** effluent concentrations of TN, $NH_4^+$-N, TP and $PO_4^{3-}$-P in Stage V; **f,** pH succession in the duplicate SPREAD UASB reactors.

**Supplementary Fig. 3 |** Excitation-emission matrix (EEM) spectra of organic matter in the SPREAD process: **a,** EEM of WAS; **b,** EEM of liquid layer organic matter; **c,** EEM of organic matter in UASB effluent; **d,** region information of the EEM spectra.

**Supplementary Fig. 4 |** Recovery efficiencies of TP and TN through struvite crystallization in SPREAD. Inside figure showed the TP/TN concentrations in the influent and effluent of the struvite crystallization step.

**Supplementary Fig. 5 |** Comparison of SPREAD UASB microbiome with conventional AD microbiomes: **a,** Principal coordinates analysis (PCoA) of microbial communities in SPREAD UASB and other digestion microbiomes (see **Supplementary Table 5** for the detailed source of other digestion microbiomes); **b,** sludge granulation in stage-V UASB reactors and floc-like sludge in conventional AD; **c,** negative/positive cohesion, **d,** sporulation score, and **e,** gene number of SPREAD UASB and conventional AD microbiomes. The influent of TAAD is the liquidized sludge without stratification.

**Supplementary Fig. 6 |** Process flowchart for construction of the anaerobic digestion database (ADDB).

**Supplementary Fig. 7 |** Phylogenetic tree of 129 MAGs retrieved from SPREAD UASB microbiome and their closely related genomes. The tree was constructed based on the concatenated alignment of 40 ribosomal proteins.

**Supplementary Fig. 8 |** Meta-omics profiles of major populations in SPREAD UASB microbiome: **a,** metagenomics potentials; **b,** metatranscriptomic profiles; **c,** metaproteomic profiles.

**Supplementary Fig. 9 |** Object and system boundary of sewage sludge anaerobic digestion processes, i.e., conventional mesophilic anaerobic digestion (CAD), anaerobic digestion combined with thermal-alkaline pretreatment (TAAD) and SPREAD

#### Supplementary Tables 1-7

**Supplementary Table 1 |** Thermodynamic data on methanogenic digestion of varied organic substrates.

**Supplementary Table 2 |** Information on metagenomics data of SPREAD UASB microbiome and other digestion microbiomes.

**Supplementary Table 3 |** Summary of MAGs retrieved from SPREAD UASB microbiome.

**Supplementary Table 4 |** Phylogeny, genome completeness, contamination and genome size of retrieved MAGs of major populations in SPREAD UASB microbiome.

**Supplementary Table 5 |** Organic carbon and energy metabolisms and associated pathways of four major populations in SPREAD UASB microbiome.

**Supplementary Table 6 |** Accession numbers of previously published sequencing data for source tracking and PCoA analysis in this study.

**Supplementary Table 7 |** Life cycle inventory of CAD, TAAD and SPREAD

#### Supplementary References

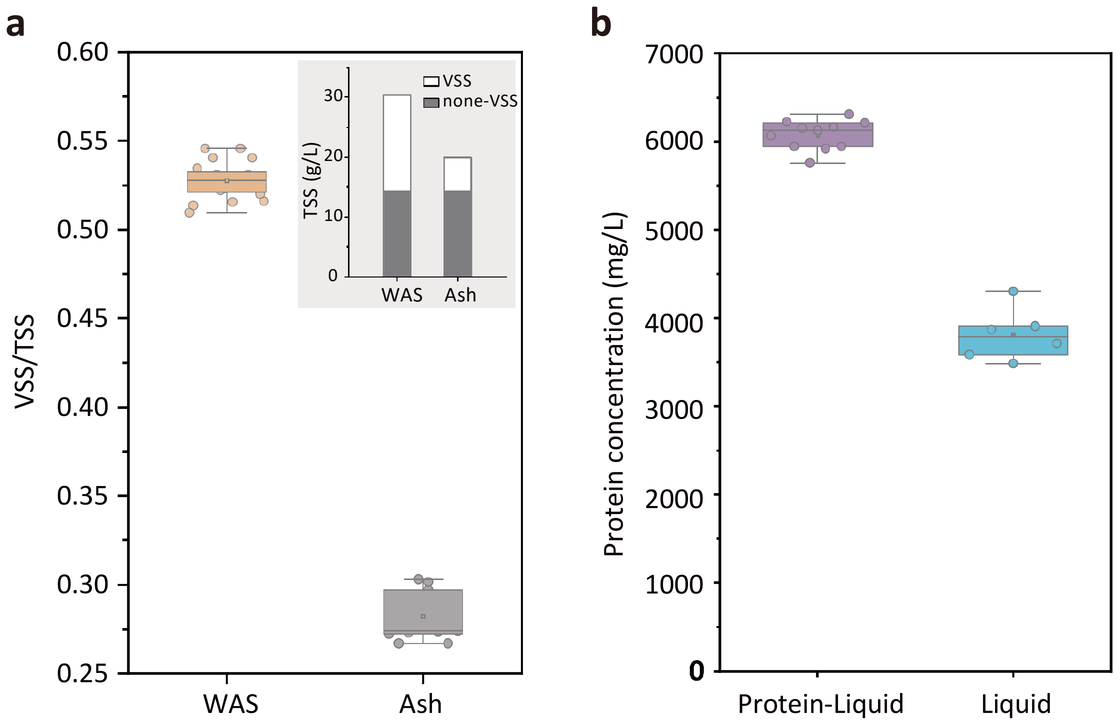


**Supplementary Fig. 1 |** Characterization of the liquefied sludge: **a,** VSS/TSS and TSS of waste activated sludge (WAS) and ash layer of the liquefied sludge; **b,** Protein concentrations in the crude-protein and liquid layers of the liquefied sludge.

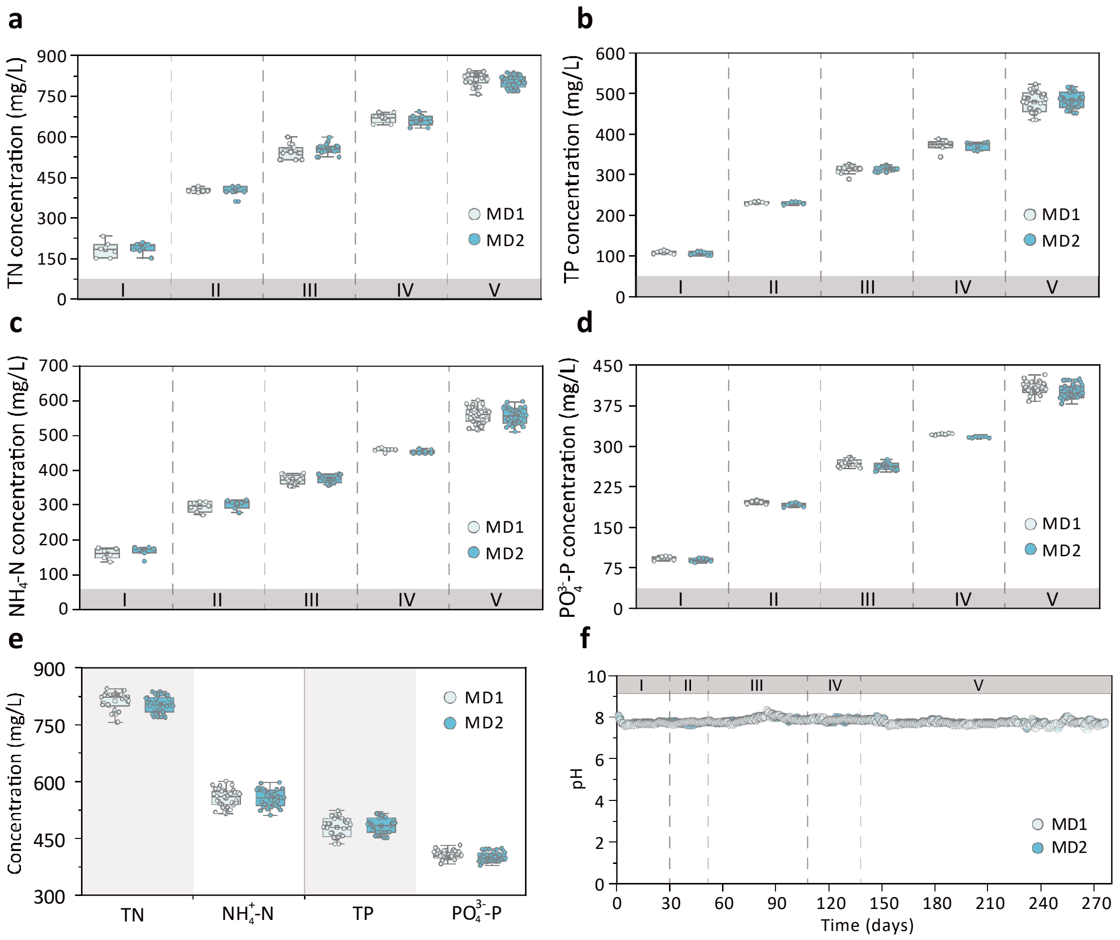


**Supplementary Fig. 2 |** Performance of the SPREAD UASB reactors MD1 and MD2: **a,** effluent TN concentrations in the five stages; **b,** effluent $NH_4^+$-N concentrations in the five stages; **c,** effluent TP concentrations in the five stages; **d,** effluent $PO_4^{3-}$-P concentrations in the five stages; **e,** effluent concentrations of TN, $NH_4^+$-N, TP and $PO_4^{3-}$-P in Stage V; **f,** pH succession in the duplicate SPREAD UASB reactors.

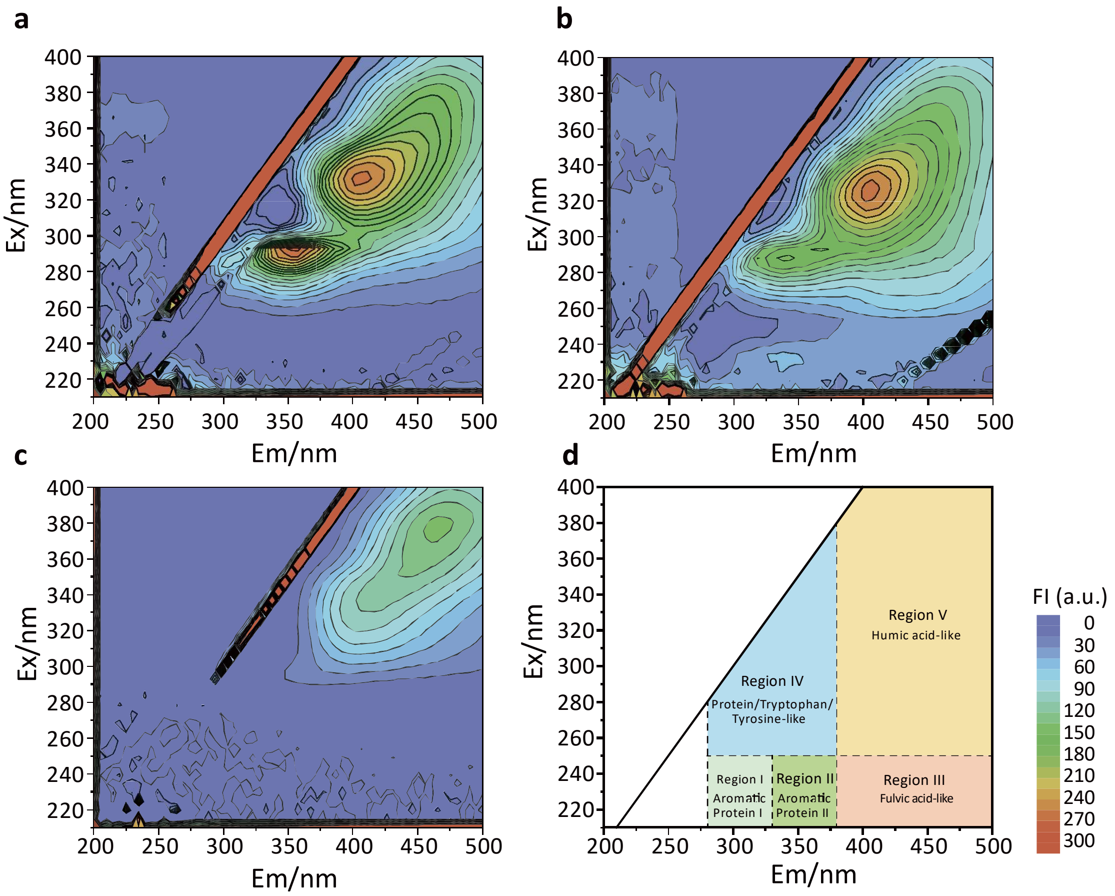


**Supplementary Fig. 3 |** Excitation-emission matrix (EEM) spectra of organic matter in the SPREAD process: **a,** EEM of WAS; **b,** EEM of liquid layer organic matter; **c,** EEM of organic matter in UASB effluent; **d,** region information of the EEM spectra.

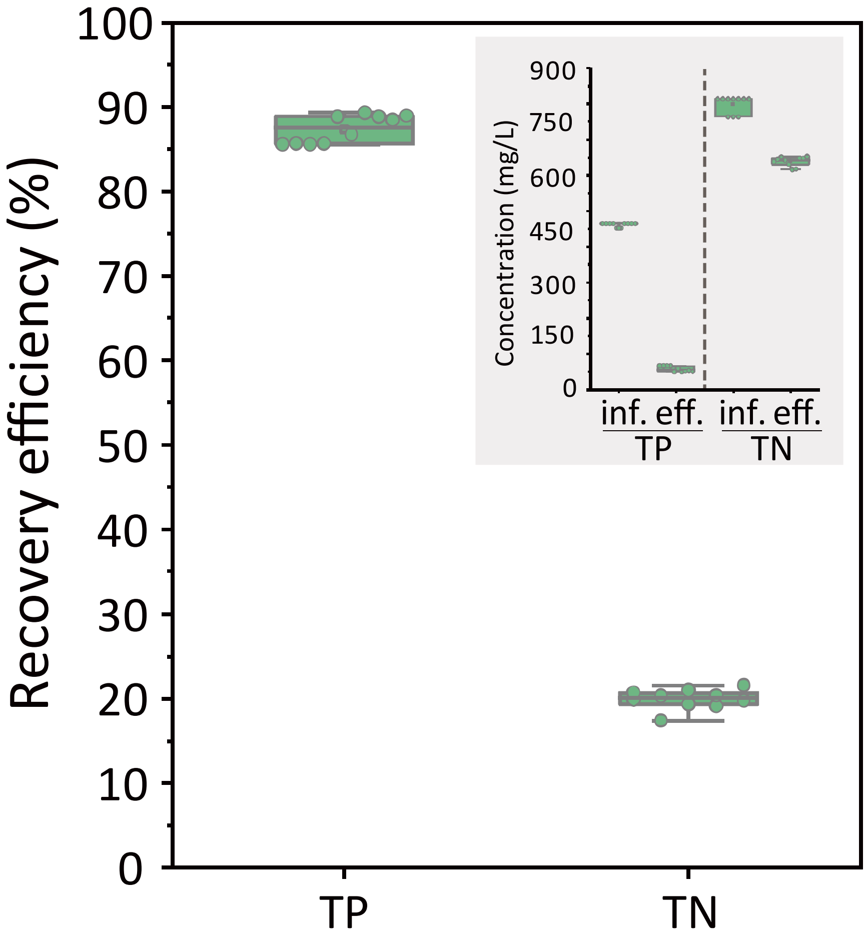


**Supplementary Fig. 4 |** Recovery efficiencies of TP and TN through struvite crystallization in SPREAD. Inside figure showed the TP/TN concentrations in the influent and effluent of the struvite crystallization step.

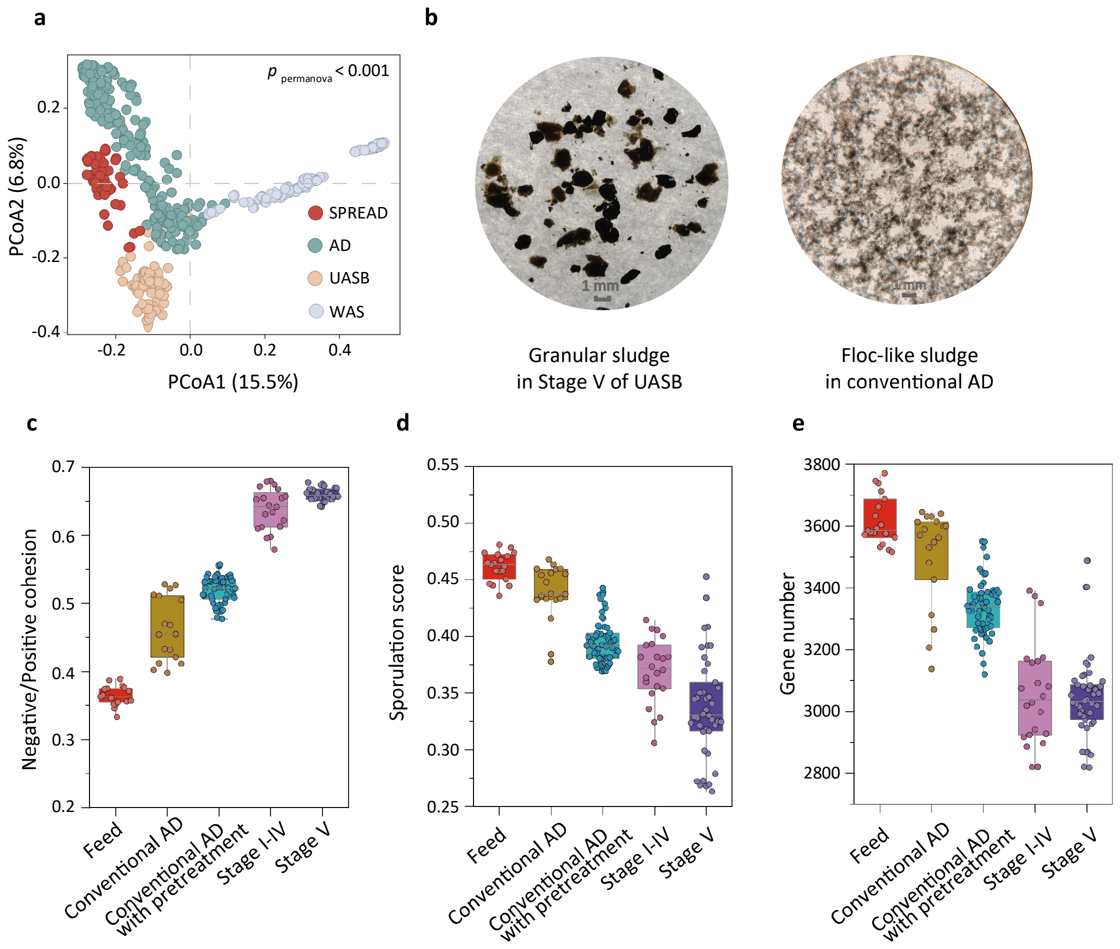


**Supplementary Fig. 5 |** Comparison of SPREAD UASB microbiome with conventional AD microbiomes: **a,** Principal coordinates analysis (PCoA) of microbial communities in SPREAD UASB and other digestion microbiomes (see **Supplementary Table 5** for the detailed source of other digestion microbiomes); **b,** sludge granulation in stage-V UASB reactors and floc-like sludge in conventional AD; **c,** negative/positive cohesion, **d,** sporulation score, and **e,** gene number of SPREAD UASB and conventional AD microbiomes. The influent of TAAD is the liquidized sludge without stratification.

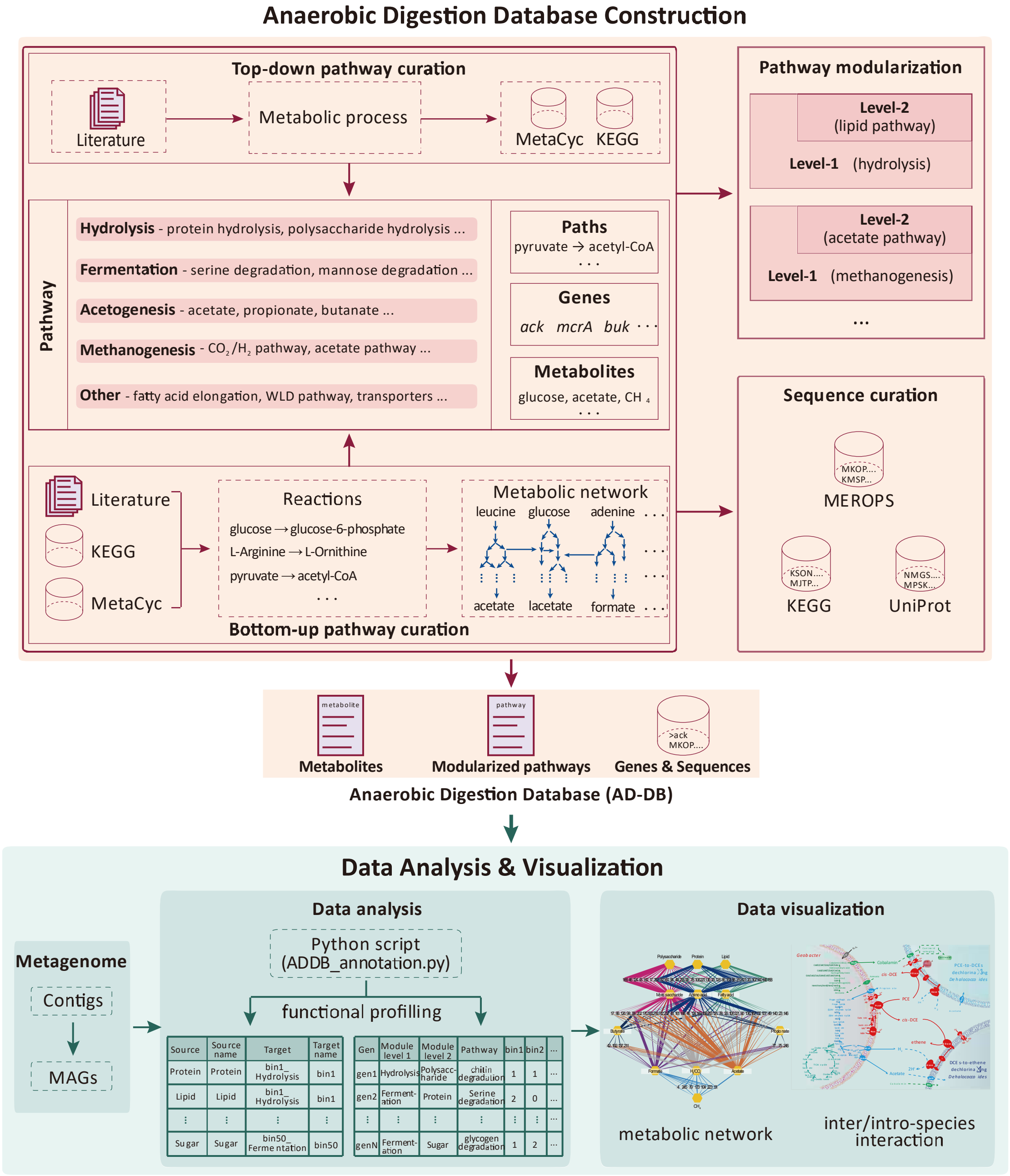


**Supplementary Fig. 6 |** Process flowchart for construction of the anaerobic digestion database (ADDB).

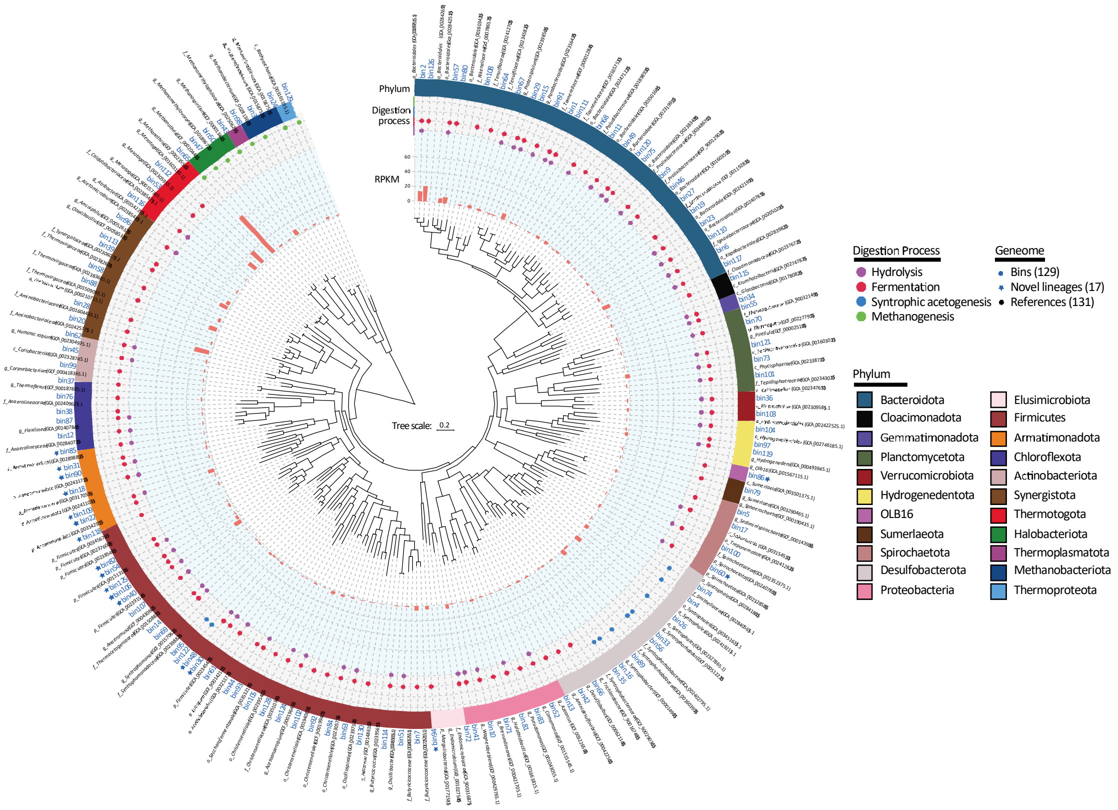


**Supplementary Fig. 7 |** Phylogenetic tree of 129 MAGs retrieved from SPREAD UASB microbiome and their closely related genomes. The tree was constructed based on the concatenated alignment of 40 ribosomal proteins.

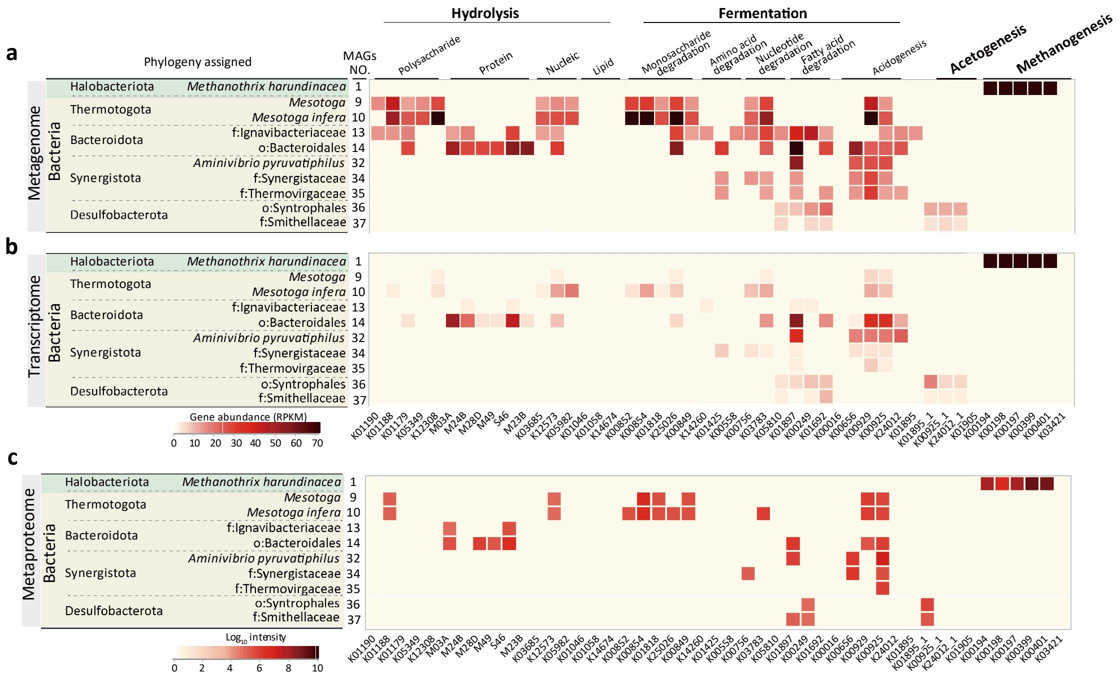


**Supplementary Fig. 8 |** Meta-omics profiles of major populations in SPREAD UASB microbiome: **a,** metagenomics potentials; **b,** metatranscriptomic profiles; **c,** metaproteomic profiles.

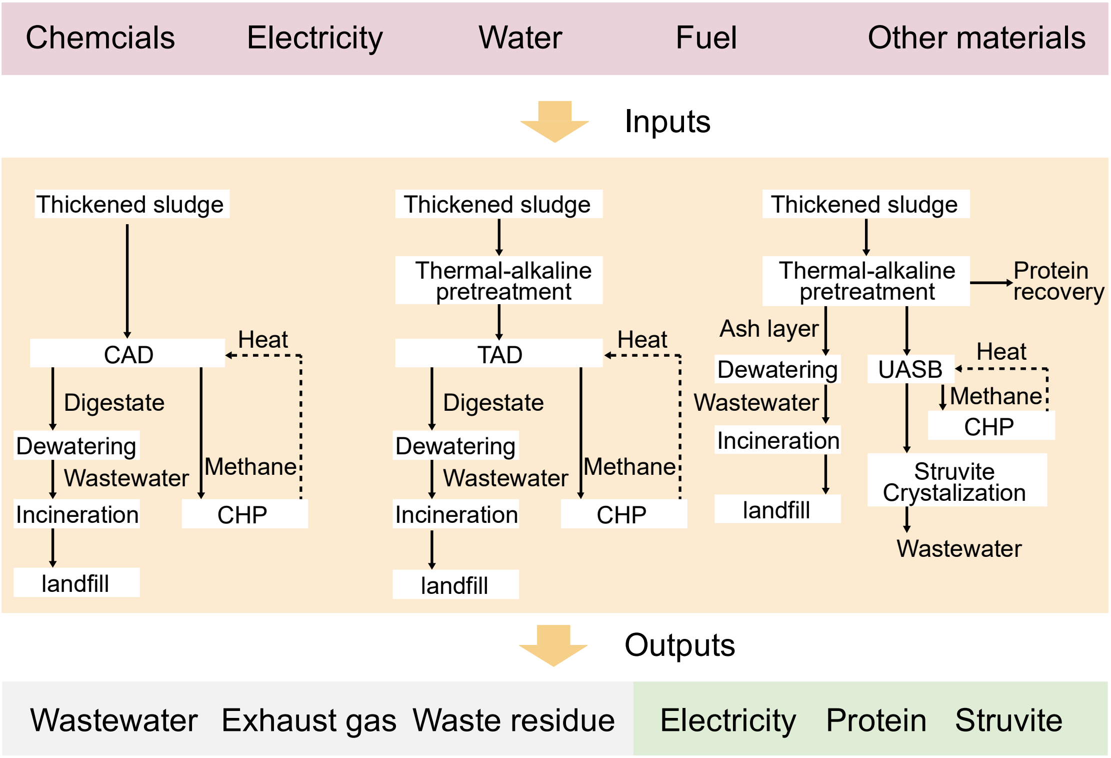


**Supplementary Fig. 9 |** System boundary of sewage sludge anaerobic digestion processes, i.e., conventional mesophilic anaerobic digestion (CAD), anaerobic digestion combined with thermal-alkaline pretreatment (TAAD) and SPREAD

**Supplementary Table 1 |** Thermodynamic data on methanogenic digestion of varied organic substrates

| Samples | Substrate | Reaction | △G' KJ / reaction | $e^-$ transfer number | △G' KJ / $e^-$ | Reference |
|---|---|---|---|---|---|---|
| DRR321573 | Terephthalic acid | $C_6H_4(COO^-)_2 + 8.75H_2O \rightarrow 3.75CH_4 + 4.25HCO_3^- + 2.25H^+$ | -107.9 | 15 | -7.19 | Kuroda et al., 2022 |
| SRR13155356<br>SRR13155357 | Acetate | $CH_3COO^- + H_2O \rightarrow CH_4 + HCO_3^-$ | -31.12 | 4 | -7.78 | Detman et al., 2021 |
| SRR10872666 | Benzoic acid | $C_6H_5COO^- + 7.75H_2O \rightarrow 3.75CH_4 + 3.25HCO_3^- + 2.25H^+$ | -118.68 | 15 | -7.92 | Gagliano et al., 2022 |
| SRR13155358<br>SRR13155359 | Propionate | $CH_3CH_2COO^- + 1.75H_2O \rightarrow 1.75CH_4 + 1.25HCO_3^- + 0.25H^+$ | -56.16 | 7 | -8.02 | Detman et al., 2021 |
| SRR13155360<br>SRR13155361 | Butyrate | $CH_3CH_2CH_2COO^- + 2.5H_2O \rightarrow 2.5CH_4 + 1.5HCO_3^- + 0.5H^+$ | -81.47 | 10 | -8.15 | Detman et al., 2021 |
| SRR13155362<br>SRR13155363 | Lactate | $CH_3CHOHCOO^- + 1.5H_2O \rightarrow 1.5CH_4 + 1.5HCO_3^- + 0.5H^+$ | -102.77 | 6 | -17.13 | Detman et al., 2021 |
| ERR1746306<br>ERR1746305 | Mixture (acetate, glucose oxytetracycline)[a] | - | -218.4 | 8 | -20.79 | Yi et al., 2017 |
| SRR16951079<br>SRR16951080 | Mixture (VFAs, maltose)[b] | - | -446.43 | 14 | -21.84 | Bovio-Winkler et al., 2023 |
| SRR7687450<br>SRR7687452<br>SRR7687449 | Mixture (VFAs, carbonhydrate)[c] | - | -446.43 | 14 | -21.84 | Lei et al., 2019 |
| SPREAD | WAS | $C_5H_7O_2N + 5.5H_2O \rightarrow 2.5CH_4 + NH_4^+ + 2.5HCO_3^- + 1.5H^+$ | -314.71 | 10 | -31.47 | This study |

**Supplementary Table 1 |** Thermodynamic data on methanogenic digestion of varied organic substrates (Cont.)

| Samples | Substrate | Reaction | △G' KJ / reaction | $e^-$ transfer number | △G' KJ / $e^-$ | Reference |
|---|---|---|---|---|---|---|
| SRR18053990<br>SRR18053991 | Black water (carbonhydrate, cells) | - | -588.23 | 17 | -33.69 | Dang et al., 2022 |
| ERR1876141 | Glucose | $C_6H_{12}O_6 + 3H_2O \rightarrow 3CH_4 + 3HCO_3^- + 3H^+$ | -405.68 | 12 | -33.81 | Jing et al., 2017 |
| SRR11058229<br>SRR11058230<br>SRR11058231 | Sugarcane | $C_{12}H_{22}O_{11} + 7H_2O \rightarrow 6CH_4 + 6HCO_3^- + 6H^+$ | -861.74 | 24 | -35.91 | Camargo et al.,2023 |
| TAAD | WAS (cells) | $C_5H_7O_2N + 5.5H_2O - > 2.5CH_4 + NH_4^+ +2.5HCO_3^- +1.5H^+$ | -314.71 | 10 | -31.47 | Wang et al., 2020 |
| APAD (ERR3454365)<br>HS-APAD (ERR3454366)<br>Thermal-AD (ERR3454364)<br>WAS-AD (ERR3454363) | WAS (cells) | $C_5H_7O_2N + 5.5H_2O - > 2.5CH_4 + NH_4^+ +2.5HCO_3^- +1.5H^+$ | -314.71 | 10 | -31.47 | Liang et al., 2020 |
| WAS-AD (ERR13624441) | WAS (cells) | $C_5H_7O_2N + 5.5H_2O - > 2.5CH_4 + NH_4^+ +2.5HCO_3^- +1.5H^+$ | -314.71 | 10 | -31.47 | Shi et al., 2022 |
| STR-FW-AD (ERR13624442)<br>STR-AD (ERR13624443)<br>FW-AD (ERR13624444) | Straw, food waste<br>Straw (carbonhydrate)<br>Food waste (carbonhydrate, protein) | $C_{12}H_{22}O_{11} + 7H_2O \rightarrow 6CH_4 + 6HCO_3^- + 6H^+$ | -861.74 | 24 | 35.91 | Shi et al., 2022 |

Note: a, the standard gibbs free energy of mixture was calculated based on 50% acetate and 50% glucose; b, the standard gibbs free energy of mixture was calculated based on 50% acetate and 50% maltose; c, the standard gibbs free energy of mixture was calculated based on 50% acetate and 50% carbonhydrate;

**Supplementary Table 2 |** Information on metagenomics data of SPREAD UASB microbiome and other digestion microbiomes.

| Item | SPREAD | SRR13155356 | SRR13155359 | SRR7687450 | ERR1876141 | SRR11058229 | TAAD | APAD | WAS |
|---|---|---|---|---|---|---|---|---|---|
| Raw reads (Gb) | **35** | 28 | 13 | 15 | 24 | 8 | 26 | 18 | 26 |
| Contigs | | | | | | | | | |
| Number of total | **1119197** | 699714 | 425801 | 443825 | 543346 | 170052 | 1527689 | 972424 | 2822034 |
| Total length | **1302774411** | 606780546 | 37084695 | 418429785 | 591247519 | 164107483 | 1138132608 | 911259186 | 1734734683 |
| Number of >=1000bp | **240263** | 110473 | 71334 | 75651 | 118493 | 29645 | 159419 | 173386 | 241243 |
| Total length of >=1000bp | **852749064** | 323078908 | 203938577 | 236234979 | 376767567 | 93009102 | 550502802 | 521688623 | 639047478 |
| Mean GC content (%) | **52.15** | 49.87 | 50.18 | 47.49 | 51.69 | 47.63 | 51.92 | 55.14 | 58.06 |
| N50 (bp) | **5785** | 3838 | 3555 | 4613 | 4402 | 4493 | 5892 | 4010 | 3163 |
| N75 (bp) | **2143** | 1769 | 1775 | 1880 | 1941 | 1767 | 1994 | 1840 | 1590 |
| Longest (bp) | **650671** | 48080 | 460665 | 252200 | 896506 | 932562 | 1516569 | 909963 | 683280 |
| Mapped read (%) | **86.92** | 62.67 | 77.34 | 77.96 | 82.81 | 84.59 | 90.12 | 85.38 | 86.66 |

**Supplementary Table 3 |** Summary of MAGs retrieved from SPREAD UASB microbiome.

| Draft genome | Completeness (%) | Contamination (%) | CDS | GC content | N50 (bp) | Size (bp) | Lineage | Abundance (RPKM) |
|---|---|---|---|---|---|---|---|---|
| | | | | | | | | SPREAD |
| bin.1.orig | 91.7 | 1.217 | 3310 | 0.456 | 5799 | 4094306 | Bacteroidota | 0.425 |
| bin.2.strict | 85.55 | 2.433 | 1783 | 0.412 | 53117 | 2043200 | Bacteroidota | 13.574 |
| bin.3.strict | 98.13 | 0 | 2145 | 0.4 | 86308 | 2183471 | Euryarchaeota | 1.583 |
| bin.4.orig | 90.99 | 3.87 | 2520 | 0.48 | 28261 | 2707391 | Desulfobacterota | 5.375 |
| bin.5.strict | 90.34 | 1.905 | 2572 | 0.52 | 10852 | 2668066 | Spirochaetota | 1.527 |
| bin.6.strict | 97.2 | 1.675 | 3135 | 0.331 | 170121 | 3784114 | Bacteroidota | 10.568 |
| bin.7.strict | 77.78 | 1.216 | 1606 | 0.575 | 7157 | 1688382 | Firmicutes_A | 5.505 |
| bin.8.orig | 82.58 | 2.247 | 1539 | 0.28 | 9601 | 1550681 | Firmicutes | 0.648 |
| bin.9.orig | 99.46 | 2.956 | 4008 | 0.475 | 251943 | 5065381 | Bacteroidota | 2.752 |
| bin.10.orig | 85.29 | 1.958 | 2532 | 0.669 | 4590 | 2619952 | Proteobacteria | 0.445 |
| bin.11.orig | 98.38 | 0.268 | 2257 | 0.344 | 205955 | 2623278 | Bacteroidota | 2.459 |
| bin.12.orig | 80.18 | 2.909 | 2195 | 0.406 | 113978 | 2478187 | Chloroflexota | 0.912 |
| bin.13.orig | 99.4 | 0.295 | 2965 | 0.669 | 28290 | 3135923 | Desulfobacterota_A | 1.034 |
| bin.14.orig | 92.86 | 1.394 | 2073 | 0.493 | 49303 | 2340587 | Firmicutes_B | 0.681 |
| bin.15.orig | 79.89 | 0.018 | 1318 | 0.364 | 21475 | 1558945 | Bacteroidota | 0.655 |
| bin.16.orig | 99.51 | 2.741 | 3537 | 0.569 | 30634 | 4068504 | Desulfobacterota | 0.959 |
| bin.17.permissive | 77.71 | 2.612 | 2405 | 0.471 | 3496 | 2572369 | Spirochaetota | 0.356 |
| bin.18.orig | 78.7 | 0 | 2372 | 0.674 | 59278 | 3054327 | Armatimonadota | 0.736 |
| bin.19.orig | 91.21 | 1.612 | 3110 | 0.469 | 75910 | 3998164 | Bacteroidota | 1.379 |
| bin.20.orig | 89.83 | 0.847 | 2105 | 0.603 | 5897 | 2214197 | Synergistota | 18.860 |
| bin.22.orig | 77.31 | 0.925 | 3141 | 0.634 | 123914 | 4080757 | Armatimonadota | 1.092 |
| bin.23.orig | 96.23 | 0.537 | 2346 | 0.456 | 99533 | 2907980 | Bacteroidota | 1.024 |
| bin.24.permissive | 91.96 | 2.97 | 2097 | 0.349 | 49606 | 1889804 | Euryarchaeota | 2.528 |

**Supplementary Table 3 |** Summary of MAGs retrieved from SPREAD UASB microbiome (Cont.)

| Draft genome | Completeness (%) | Contamination (%) | CDS | GC content | N50 (bp) | Size (bp) | Lineage | Abundance (RPKM) |
|---|---|---|---|---|---|---|---|---|
| | | | | | | | | SPREAD |
| bin.25.orig | 71.41 | 1.907 | 1568 | 0.553 | 4065 | 1613143 | Synergistota | 1.786 |
| bin.26.orig | 85.8 | 2.58 | 2242 | 0.639 | 46221 | 2389345 | Desulfobacterota | 8.581 |
| bin.27.orig | 98.57 | 3.809 | 2606 | 0.32 | 48640 | 3252213 | Bacteroidota | 1.060 |
| bin.28.strict | 93.22 | 2.526 | 3573 | 0.591 | 13305 | 3847128 | Synergistota | 1.050 |
| bin.29.orig | 83.6 | 1.092 | 1898 | 0.46 | 19343 | 2290301 | Bacteroidota | 0.737 |
| bin.30.strict | 93.14 | 0.211 | 2175 | 0.496 | 29401 | 2308958 | Firmicutes_D | 2.246 |
| bin.31.orig | 96.29 | 0 | 2929 | 0.549 | 72642 | 3537525 | Armatimonadota | 0.715 |
| bin.33.orig | 93.27 | 3.781 | 2605 | 0.52 | 28477 | 2735271 | Desulfobacterota | 1.959 |
| bin.34.orig | 97.8 | 0 | 3436 | 0.603 | 32486 | 4376645 | Gemmatimonadota | 0.780 |
| bin.35.strict | 84.32 | 1.29 | 3128 | 0.586 | 7440 | 3318895 | Desulfobacterota | 0.667 |
| bin.36.orig | 91.04 | 1.718 | 2045 | 0.615 | 18363 | 2374073 | Verrucomicrobiota | 0.658 |
| bin.37.orig | 79.78 | 2.935 | 2758 | 0.665 | 10563 | 3119342 | Chloroflexota | 0.525 |
| bin.38.orig | 77.36 | 2.272 | 2138 | 0.554 | 5517 | 2217193 | Chloroflexota | 0.410 |
| bin.39.strict | 94.6 | 0.308 | 1458 | 0.523 | 54215 | 1590503 | Synergistota | 10.729 |
| bin.40.strict | 78.21 | 0 | 1236 | 0.469 | 29375 | 1364093 | Firmicutes_E | 9.217 |
| bin.41.orig | 88.97 | 1.477 | 3346 | 0.662 | 6838 | 3595383 | Proteobacteria | 0.476 |
| bin.42.orig | 97.71 | 0.892 | 2935 | 0.628 | 22516 | 3383151 | Desulfobacterota | 1.896 |
| bin.43.permissive | 94.39 | 0 | 2038 | 0.458 | 33846 | 2134979 | Halobacterota | 8.254 |
| bin.44.orig | 78.31 | 1.235 | 2445 | 0.428 | 5569 | 2937495 | Firmicutes_A | 0.426 |
| bin.45.strict | 97.08 | 1.944 | 1730 | 0.661 | 45827 | 1810464 | Actinobacteriota | 0.770 |
| bin.46.permissive | 91.46 | 1.72 | 1993 | 0.423 | 18120 | 2366335 | Bacteroidota | 0.800 |
| bin.47.orig | 98.03 | 0.653 | 2168 | 0.542 | 60744 | 2205102 | Halobacterota | 1.605 |
| bin.48.strict | 91.94 | 2.817 | 2075 | 0.441 | 95313 | 2298960 | Firmicutes_D | 0.878 |

**Supplementary Table 3 |** Summary of MAGs retrieved from SPREAD UASB microbiome (Cont.)

| Draft genome | Completeness (%) | Contamination (%) | CDS | GC content | N50 (bp) | Size (bp) | Lineage | Abundance (RPKM) |
|---|---|---|---|---|---|---|---|---|
| | | | | | | | | SPREAD |
| bin.49.orig | 96.74 | 3.333 | 2940 | 0.451 | 195089 | 3712917 | Bacteroidota | 3.522 |
| bin.50.strict | 91.02 | 2.339 | 1864 | 0.429 | 4244 | 1829765 | Halobacterota | 0.391 |
| bin.51.orig | 91.52 | 1.363 | 2545 | 0.605 | 13747 | 2754742 | Firmicutes_A | 0.629 |
| bin.52.orig | 93.5 | 1.086 | 3845 | 0.625 | 16983 | 4161595 | Proteobacteria | 0.564 |
| bin.53.strict | 97.3 | 2.664 | 2531 | 0.485 | 83099 | 2812049 | Thermotogota | 12.066 |
| bin.54.orig | 90.59 | 0 | 2241 | 0.56 | 172944 | 2455950 | Firmicutes_E | 2.292 |
| bin.55.orig | 88 | 2.612 | 4190 | 0.544 | 5026 | 4955066 | Planctomycetota | 0.448 |
| bin.56.orig | 75.08 | 4.806 | 1996 | 0.439 | 3913 | 2137194 | Desulfobacterota | 7.202 |
| bin.57.orig | 90.31 | 0.476 | 1636 | 0.405 | 19358 | 1911247 | Bacteroidota | 5.500 |
| bin.58.orig | 83.05 | 0 | 1532 | 0.541 | 13784 | 1586656 | Synergistota | 3.506 |
| bin.59.strict | 73.44 | 3.318 | 2794 | 0.672 | 4270 | 2824029 | Pseudomonadota | 1.931 |
| bin.60.strict | 91.31 | 1.123 | 2910 | 0.575 | 54250 | 3268827 | Spirochaetota | 2.012 |
| bin.61.orig | 91.61 | 3.629 | 3135 | 0.403 | 18414 | 3424807 | Firmicutes_A | 0.644 |
| bin.62.orig | 97.09 | 0.909 | 1516 | 0.347 | 43624 | 1647858 | Actinobacteriota | 0.947 |
| bin.63.orig | 93.62 | 2.348 | 2235 | 0.552 | 12852 | 2673241 | Firmicutes_A | 0.565 |
| bin.64.orig | 98.57 | 2.777 | 3076 | 0.429 | 70059 | 3755761 | Bacteroidota | 3.830 |
| bin.65.strict | 98.68 | 0 | 2065 | 0.625 | 31450 | 2050991 | Halobacterota | 64.970 |
| bin.66.orig | 97.41 | 0.645 | 3176 | 0.534 | 197301 | 3410355 | Desulfuromonadota | 1.489 |
| bin.67.orig | 92.85 | 1.428 | 2357 | 0.443 | 84002 | 2983603 | Bacteroidota | 2.042 |
| bin.68.orig | 97.58 | 1.075 | 2462 | 0.434 | 40328 | 2978624 | Bacteroidota | 0.735 |
| bin.69.orig | 100 | 1.858 | 3042 | 0.43 | 78962 | 3199657 | Firmicutes_B | 0.899 |
| bin.70.orig | 93.1 | 0 | 3462 | 0.623 | 30026 | 4503583 | Planctomycetota | 0.626 |
| bin.71.orig | 97.38 | 0.095 | 2547 | 0.631 | 29017 | 2782290 | Proteobacteria | 2.613 |

**Supplementary Table 3 |** Summary of MAGs retrieved from SPREAD UASB microbiome (Cont.)

| Draft genome | Completeness (%) | Contamination (%) | CDS | GC content | N50 (bp) | Size (bp) | Lineage | Abundance (RPKM) |
|---|---|---|---|---|---|---|---|---|
| | | | | | | | | SPREAD |
| bin.72.strict | 97.75 | 1.123 | 1593 | 0.346 | 69540 | 1881903 | Elusimicrobiota | 5.961 |
| bin.73.orig | 100 | 2.272 | 4166 | 0.626 | 161712 | 5360700 | Planctomycetota | 5.064 |
| bin.74.orig | 87.09 | 1.935 | 2822 | 0.618 | 48455 | 3149268 | Desulfobacterota | 0.720 |
| bin.75.orig | 99.52 | 2.619 | 2875 | 0.469 | 290074 | 3518095 | Bacteroidota | 2.100 |
| bin.76.strict | 79.09 | 0.909 | 1685 | 0.505 | 73413 | 1884462 | Chloroflexota | 1.865 |
| bin.77.strict | 73 | 2.331 | 0.608 | 3050 | 4534 | 3229900 | Firmicutes_A | 1.482 |
| bin.78.orig | 72.21 | 3.448 | 0.458 | 2665 | 6670 | 3308408 | Firmicutes_A | 2.315 |
| bin.79.strict | 89.51 | 1.648 | 2667 | 0.596 | 17192 | 3246612 | Sumerlaeota | 0.616 |
| bin.80.strict | 93.49 | 2.539 | 1851 | 0.447 | 54530 | 2158758 | Bacteroidota | 7.708 |
| bin.81.orig | 85.2 | 1.534 | 3452 | 0.564 | 6999 | 3611938 | Proteobacteria | 0.459 |
| bin.82.strict | 87.62 | 0 | 2331 | 0.572 | 55114 | 2520664 | Firmicutes_E | 0.773 |
| bin.83.strict | 92.29 | 2.207 | 2311 | 0.609 | 34887 | 2603903 | Proteobacteria | 1.062 |
| bin.84.orig | 93.11 | 1.612 | 2231 | 0.62 | 16150 | 2380012 | Firmicutes_A | 0.579 |
| bin.85.orig | 82.71 | 1.028 | 2869 | 0.61 | 4129 | 3385913 | Armatimonadota | 0.383 |
| bin.86.strict | 98.83 | 3.296 | 3900 | 0.517 | 70487 | 4980742 | OLB16 | 0.781 |
| bin.87.strict | 82.12 | 0 | 1882 | 0.474 | 32606 | 2131275 | Chloroflexota | 0.528 |
| bin.88.orig | 100 | 0 | 1696 | 0.566 | 46137 | 1758198 | Synergistota | 10.238 |
| bin.89.orig | 94.19 | 1.29 | 3162 | 0.573 | 82963 | 3373276 | Desulfobacterota | 1.353 |
| bin.90.orig | 95.37 | 0 | 2623 | 0.53 | 180581 | 3082903 | Armatimonadota | 1.487 |
| bin.91.strict | 81.28 | 4.184 | 2046 | 0.45 | 12940 | 2339254 | Bacteroidota | 7.962 |
| bin.92.orig | 91.67 | 2.622 | 3343 | 0.61 | 15288 | 3896622 | Firmicutes_A | 0.682 |
| bin.93.permissive | 84.21 | 4.086 | 2436 | 0.613 | 8974 | 2758998 | Firmicutes_A | 0.700 |
| bin.94.strict | 76.21 | 1.139 | 1013 | 0.512 | 16379 | 1024165 | Margulisbacteria | 3.957 |

**Supplementary Table 3 |** Summary of MAGs retrieved from SPREAD UASB microbiome (Cont.)

| Draft genome | Completeness (%) | Contamination (%) | CDS | GC content | N50 (bp) | Size (bp) | Lineage | Abundance (RPKM) |
|---|---|---|---|---|---|---|---|---|
| | | | | | | | | SPREAD |
| bin.95.orig | 96.28 | 2.933 | 2260 | 0.5 | 31359 | 2406223 | Firmicutes_B | 2.093 |
| bin.96.strict | 75.14 | 0.847 | 2383 | 0.611 | 35235 | 2781155 | Synergistota | 0.829 |
| bin.97.orig | 98.9 | 2.197 | 4431 | 0.659 | 91355 | 5718988 | Hydrogenedentota | 1.344 |
| bin.98.strict | 96.37 | 0 | 1239 | 0.536 | 261291 | 1179858 | Thermoplasmatota | 0.952 |
| bin.99.orig | 98.87 | 0.563 | 2126 | 0.666 | 88748 | 2356782 | Actinobacteriota | 0.834 |
| bin.100.orig | 91.95 | 1.149 | 2755 | 0.663 | 4016 | 3001730 | Spirochaetota | 0.434 |
| bin.101.orig | 96.3 | 3.512 | 4199 | 0.644 | 22074 | 5292798 | Planctomycetota | 0.600 |
| bin.102.orig | 88.3 | 0 | 2320 | 0.551 | 115451 | 2474175 | Firmicutes_A | 2.272 |
| bin.103.orig | 94.59 | 3.407 | 2971 | 0.648 | 107921 | 4144021 | Verrucomicrobiota | 1.136 |
| bin.104.orig | 90.04 | 4.395 | 3500 | 0.632 | 8968 | 4317129 | Hydrogenedentota | 0.457 |
| bin.105.orig | 96.34 | 0 | 1787 | 0.392 | 130516 | 1955783 | Firmicutes_A | 1.341 |
| bin.106.orig | 91.08 | 0 | 2083 | 0.49 | 143251 | 2352379 | Firmicutes_E | 2.641 |
| bin.107.strict | 87.23 | 3.571 | 2489 | 0.504 | 7312 | 2614542 | Firmicutes_C | 0.423 |
| bin.108.strict | 91.82 | 0.961 | 1720 | 0.553 | 102603 | 2135972 | Bacteroidota | 1.678 |
| bin.109.strict | 93.51 | 4.506 | 5679 | 0.678 | 93938 | 8621854 | Armatimonadota | 7.500 |
| bin.110.orig | 98.11 | 0.537 | 1879 | 0.331 | 62726 | 2237594 | Bacteroidota | 2.225 |
| bin.111.orig | 97.69 | 0.769 | 3001 | 0.489 | 39656 | 3660692 | Bacteroidota | 0.802 |
| bin.112.permissive | 98.11 | 0 | 2574 | 0.504 | 94202 | 2911912 | Thermotogota | 19.725 |
| bin.113.orig | 94.23 | 0 | 1780 | 0.483 | 45210 | 1951455 | Synergistota | 4.840 |
| bin.114.strict | 76.62 | 0.915 | 1593 | 0.588 | 5857 | 1717621 | Firmicutes_A | 0.534 |
| bin.115.orig | 97.8 | 0 | 1717 | 0.505 | 61589 | 2234297 | Cloacimonadota | 1.501 |
| bin.116.permissive | 91.37 | 2.033 | 1837 | 0.39 | 4269 | 2094660 | Caldatribacteriota | 0.434 |
| bin.117.strict | 97.97 | 0.546 | 2018 | 0.394 | 53700 | 2651645 | Bacteroidota | 0.776 |

**Supplementary Table 3 |** Summary of MAGs retrieved from SPREAD UASB microbiome (Cont.)

| Draft genome | Completeness (%) | Contamination (%) | CDS | GC content | N50 (bp) | Size (bp) | Lineage | Abundance (RPKM) |
|---|---|---|---|---|---|---|---|---|
| | | | | | | | | SPREAD |
| bin.118.strict | 94.23 | 0.48 | 1994 | 0.411 | 78990 | 2256813 | Firmicutes_G | 0.960 |
| bin.119.orig | 96.77 | 2.15 | 2540 | 0.526 | 62492 | 3335433 | Hydrogenedentota | 3.159 |
| bin.120.orig | 88.76 | 3.269 | 2640 | 0.456 | 9131 | 3242360 | Bacteroidota | 0.448 |
| bin.121.orig | 93.03 | 0 | 3978 | 0.668 | 16651 | 5119772 | Planctomycetota | 0.566 |
| bin.122.strict | 91.09 | 1.569 | 1932 | 0.494 | 22854 | 2047323 | Firmicutes_B | 0.839 |
| bin.123.strict | 71.22 | 2.37 | 3165 | 0.647 | 3699 | 3664844 | Hydrogenedentota | 0.331 |
| bin.124.strict | 83.28 | 1.631 | 1938 | 0.333 | 9076 | 2090324 | Firmicutes_A | 3.942 |
| bin.125.orig | 82.15 | 0 | 1750 | 0.483 | 6309 | 1976784 | Firmicutes_E | 0.416 |
| bin.126.orig | 92.69 | 1.19 | 1839 | 0.4 | 72057 | 2096081 | Bacteroidota | 20.720 |
| bin.127.orig | 70.11 | 1.151 | 4403 | 0.651 | 4897 | 4740077 | Bacteroidota | 0.878 |
| bin.128.permissive | 76.63 | 1.612 | 1614 | 0.525 | 3988 | 1653979 | Firmicutes_A | 0.396 |
| bin.129.permissive | 81.46 | 4.205 | 1288 | 0.381 | 3745 | 1281196 | Crenarchaeota | 0.369 |
| bin.130.strict | 75.34 | 2.516 | 2018 | 0.504 | 4034 | 2169996 | Firmicutes_A | 0.396 |
| bin.131.strict | 72.02 | 2.013 | 2537 | 0.535 | 3847 | 2737868 | Firmicutes_A | 0.376 |

**Supplementary Table 4 |** Phylogeny, genome completeness, contamination and genome size of retrieved MAGs of major populations in SPREAD UASB microbiome.

| MAG_NO. | Completeness | Contamination | GC content | Phylum | N50 | size (Mb) |
|---|---|---|---|---|---|---|
| 1 | 98.68 | 0 | 0.625 | Halobacteriota | 31450 | 2.050991 |
| 2 | 96.4 | 0.653 | 0.526 | Halobacteriota | 23278 | 2.242103 |
| 3 | 95.86 | 1.307 | 0.551 | Halobacteriota | 19017 | 1.688952 |
| 4 | 97.35 | 0.653 | 0.423 | Halobacteriota | 10977 | 3.171201 |
| 5 | 97.98 | 0 | 0.551 | Thermoplasmatota | 110894 | 1.530559 |
| 6 | 93.33 | 3.36 | 0.367 | Methanobacteriota | 13044 | 1.827319 |
| 7 | 90.51 | 2.068 | 0.31 | Thermotogota | 30558 | 1.923779 |
| 8 | 89.23 | 0 | 0.399 | Thermotogota | 17822 | 1.72293 |
| 9 | 99.52 | 0.47 | 0.485 | Thermotogota | 37545 | 2.617281 |
| 10 | 99.84 | 0 | 0.503 | Thermotogota | 82481 | 2.611732 |
| 11 | 97.9 | 1.111 | 0.373 | Bacteroidota | 34671 | 2.901194 |
| 12 | 94.98 | 1.075 | 0.466 | Bacteroidota | 50444 | 4.222915 |
| 13 | 97.2 | 1.675 | 0.331 | Bacteroidota | 170121 | 3.784114 |
| 14 | 92.69 | 1.19 | 0.4 | Bacteroidota | 72057 | 2.096081 |
| 15 | 95.14 | 1.19 | 0.461 | Bacteroidota | 251589 | 2.013365 |
| 16 | 95.43 | 0.537 | 0.376 | Bacteroidota | 79712 | 2.605574 |
| 17 | 83.76 | 1.075 | 0.351 | Bacteroidota | 28059 | 1.74912 |
| 18 | 100 | 0.54 | 0.34 | Bacteroidota | 151288 | 3.00201 |
| 19 | 98.51 | 0.787 | 0.45 | Proteobacteria | 657457 | 2.618417 |
| 20 | 99.11 | 0.662 | 0.655 | Actinobacteriota | 311048 | 3.145784 |
| 21 | 97.08 | 1.666 | 0.658 | Actinobacteriota | 1114606 | 1.916116 |
| 22 | 98.18 | 6.06 | 0.502 | Chloroflexota | 88069 | 3.342362 |
| 23 | 84.54 | 2.909 | 0.631 | Chloroflexota | 35360 | 3.988911 |

**Supplementary Table 4 |** Phylogeny, genome completeness, contamination and genome size of retrieved MAGs of major populations in SPREAD UASB microbiome (Cont)

| MAG_NO. | Completeness | Contamination | GC content | Phylum | N50 | size (Mb) |
|---|---|---|---|---|---|---|
| 24 | 84.39 | 3.212 | 0.533 | Chloroflexota | 18666 | 1.976846 |
| 25 | 98.87 | 0 | 0.442 | Firmicutes | 927734 | 1.295733 |
| 26 | 98.87 | 0 | 0.453 | Firmicutes | 311678 | 1.219889 |
| 27 | 97.17 | 1.209 | 0.605 | Bacillota | 264215 | 2.552138 |
| 28 | 85.71 | 1.785 | 0.448 | Coprothermobacterota | 14795 | 1.008339 |
| 29 | 95.9 | 0 | 0.459 | Spirochaetota | 26336 | 2.232001 |
| 30 | 98.85 | 0 | 0.571 | Spirochaetota | 34881 | 2.543419 |
| 31 | 94.29 | 0 | 0.404 | Ratteibacteria | 291813 | 1.181493 |
| 32 | 89.83 | 0.847 | 0.603 | Synergistota | 5897 | 2.214197 |
| 33 | 96.61 | 0 | 0.416 | Synergistota | 80724 | 2.268103 |
| 34 | 94.6 | 0.308 | 0.523 | Synergistota | 54215 | 1.590503 |
| 35 | 100 | 0 | 0.563 | Synergistota | 137589 | 1.878039 |
| 36 | 85.8 | 2.58 | 0.639 | Desulfobacterota | 46221 | 2.389345 |
| 37 | 90.99 | 3.87 | 0.48 | Desulfobacterota | 28261 | 2.707391 |

**Supplementary Table 5 |** Organic carbon and energy metabolisms and associated pathways of four major populations in SPREAD UASB microbiome.

| Pathway and related metabolism | Name | Enzymes | EC number | Draft genome: MAG.10 (Mesotoga) | MAG. 14 (Bacteroidales) | MAG.1 (Methanothrix) | MAG. 36 (Synrophales) |
|---|---|---|---|---|---|---|---|
| Glycolysis pathway — Embden-Meyerhof-Parnas (EMP) | glk | Glucokinase | EC:2.7.1.2 | 2 | 2 | 1 | 1 |
| | pgm | Phosphoglucomutase | EC 5.4.2.2 | 2 | 1 | | 1 |
| | pgi | Glucose-6-phosphate isomerase (phosphoglucose isomerases) | EC 5.3.1.9 | 1 | 1 | 1 | 1 |
| | pfk | ATP-dependent phosphofructokinase (PFK-A) | EC 2.7.1.11 | 3 | 1 | 1 | 1 |
| | fba | Fructose-bisphosphate aldolase | EC 4.1.2.13 | 2 | 1 | | 1 |
| | tpi | Triosephosphate isomerase | EC 5.3.1.1 | 1 | 1 | 1 | 1 |
| | gapA | Glyceraldehyde 3-phosphate dehydrogenase | EC 1.2.1.12 | 1 | 1 | | 1 |
| | pgk | Phosphoglycerate kinase | EC 2.7.2.3 | 1 | 1 | 1 | 1 |
| | eno | Enolase | EC 4.2.1.11 | 1 | 1 | 1 | 1 |
| | pyk | Pyruvate kinase | EC 2.7.1.40 | 2 | 1 | | 1 |
| Pentose phosphate pathway | tkt | Transketolase | EC:2.2.1.1 | 1 | 2 | | 1 |
| | tal | Tansaldolase | EC:2.2.1.2 | 2 | | | 1 |
| | prsA | Ribose-phosphate pyrophosphokinase | EC:2.7.6.1 | 1 | 1 | | 1 |
| | rpiB | Ribose 5-phosphate isomerase B | EC:5.3.1.6 | 1 | 1 | | 1 |
| | rpe | Ribulose-phosphate 3-epimerase | EC:5.1.3.1 | 1 | 1 | | |
| Sucrose degradation | malZ | Alpha-glucosidase | EC:3.2.1.20 | 1 | | | |
| | scrK | Fructokinase | EC:2.7.1.4 | 2 | | | |
| Xylose degradation | xylA | Xylose isomeras | EC:5.3.1.5 | 1 | | | |
| | xylB | D-xylose 1-dehydrogenase | EC:1.1.1.175 | 1 | | | |
| Wood-Ljungdahl pathway | Fhs | Formate--tetrahydrofolate ligase | [EC:6.3.4.3 | 1 | 1 | | 1 |
| | folD | Methylenetetrahydrofolate dehydrogenase (NADP+) / methenyltetrahydrofolate cyclohydrolase | EC:1.5.1.5 | 1 | 1 | 1 | |
| | metF | Methylenetetrahydrofolate reductase (NADH) | EC:1.5.1.54 | | 1 | | |
| | cdhE | Acetyl-CoA decarbonylase/synthase | EC:2.1.1.245 | | | 1 | |
| Glycine degradation | ltaE | Threonine aldolase | EC:4.1.2.48 | 1 | | | 1 |
| | glyA | Glycine hydroxymethyltransferase | EC:2.1.2.1 | 1 | 1 | 1 | 1 |
| | sda | L-serine dehydratase | EC:4.3.1.17 | 2 | 1 | | |
| | cysK | Cysteine synthase | EC:2.5.1.47 | 1 | | | |
| | cysE | Serine O-acetyltransferase | EC:2.3.1.30 | 1 | | | |
| Cysteine degradation | patB | Cysteine-S-conjugate beta-lyase | EC:4.4.1.13 | 2 | 1 | | |
| | racX | Amino-acid racemase | EC:5.1.1.10 | 1 | 1 | | |
| | dcyD | D-cysteine desulfhydrase | EC:4.4.1.15 | 1 | 1 | | |
| | aspC | Aspartate aminotransferase | EC:2.6.1.1 | | 1 | 1 | |
| | TST | Thiosulfate/3-mercaptopyruvate sulfurtransferase | EC:2.8.1.1 | | 1 | 1 | 1 |

**Supplementary Table 5 |** Organic carbon and energy metabolisms and associated pathways of four major populations in SPREAD UASB microbiome (Cont.)

| **Pathway and related metabolism** | | **Name** | **Enzymes** | **EC number** | **Draft genome** | | | |
|---|---|---|---|---|---|---|---|---|
| | | | | | **MAG.10 (Mesotoga)** | **MAG. 14 (Bacteroidales)** | **MAG.1 (Methanothrix)** | **MAG. 36 (Synrophales)** |
| Lysine degradation | | racX | Amino-acid racemase | EC:5.1.1.10 | 1 | 1 | | |
| | | kamA | Lysine 2,3-aminomutase | EC:5.4.3.3 | 1 | 2 | 2 | 3 |
| | | kamD | Beta-lysine 5,6-aminomutase alpha subunit | EC:5.4.3.3 | 1 | 1 | | |
| | | kamE | Beta-lysine 5,6-aminomutase beta subunit | EC:5.4.3.3 | 1 | 1 | | |
| | | kdd | L-erythro-3,5-diaminohexanoate dehydrogenase | EC:1.4.1.11 | 1 | 2 | | |
| | | kce | 3-keto-5-aminohexanoate cleavage enzyme | EC:2.3.1.247 | 1 | 1 | | 1 |
| | | kal | 3-aminobutyryl-CoA ammonia-lyase | EC:4.3.1.14 | 1 | 1 | | 1 |
| Histidine degradation | | hutH | Histidine ammonia-lyase | EC:4.3.1.3 | 1 | 2 | | |
| | | hutU | Urocanate hydratase | EC:4.2.1.49 | 1 | 1 | | |
| | | hutI | Imidazolonepropionase | EC:3.5.2.7 | 1 | 1 | | |
| | | fctD | Glutamate formiminotransferase / 5-formyltetrahydrofolate cyclo-ligase | EC:2.1.2.5 | 1 | 1 | | 1 |
| Glutamate | | glnA | Glutamine synthetase | EC:6.3.1.2 | 1 | 2 | 2 | 1 |
| | | gltD | Glutamate synthase (NADPH) small chain | EC:1.4.1.13 | 1 | 2 | 1 | 1 |
| | | GLUD1 | Glutamate dehydrogenase | EC:1.4.1.4 | 1 | 1 | 1 | 1 |
| Aspartate | | purA | Adenylosuccinate synthase | EC:6.3.4.4 | 1 | 1 | | 1 |
| | | purB | Adenylosuccinate lyase | EC:4.3.2.2 | 1 | 1 | 1 | 1 |
| | | argG | Argininosuccinate synthase | EC:6.3.4.5 | | 1 | 1 | 1 |
| | | argH | Argininosuccinate lyase | EC:4.3.2.1 | | 1 | 1 | 1 |
| | | arcA | Arginine deiminase | EC:3.5.3.6 | | 1 | | |
| | | asnB | Asparagine synthase (glutamine-hydrolysing) | EC:6.3.5.4 | 1 | 1 | 1 | |
| | | NIT2 | Omega-amidase | EC:3.5.1.3 | 1 | 1 | | |
| | | nadB | L-aspartate oxidase | EC:1.4.3.16 | | 1 | | 1 |
| Fermentation | Acetate fermentation | acdAB | Acetate--CoA ligase | EC:6.2.1.13 | | | 1 | 4 |
| | | ackA | Acetate kinase | EC:2.7.2.1 | 1 | 1 | | 1 |
| | | acs | Acetyl-CoA synthetase | EC 6.2.1.1 | | | 4 | |
| | Butanoate fermentation | ACAT | Acetyl-CoA C-acetyltransferase | EC:2.3.1.9 | 1 | 1 | 1 | 1 |
| | | paaH | 3-hydroxybutyryl-CoA dehydrogenase | EC:1.1.1.157 | | 1 | | 2 |
| | | crt | Enoyl-CoA hydratase | EC:4.2.1.17 | | 1 | | 3 |
| | | croR | 3-hydroxybutyryl-CoA dehydratase | EC:4.2.1.55 | 1 | 1 | | 1 |
| | | ccrA | Crotonyl-CoA reductase | EC:1.3.1.86 | | | | |
| | | ptb | Phosphate butyryltransferase | EC:2.3.1.19 | 2 | 1 | | 2 |
| | | buk | Butyrate kinase | EC:2.7.2.7 | 3 | 1 | | |
| | | atoA | Acetate CoA/acetoacetate CoA-transferase alpha subunit | EC:2.8.3.8 | 1 | 1 | | |
| | | atoD | Acetate CoA/acetoacetate CoA-transferase alpha subunit | EC:2.8.3.8 | 1 | 1 | | |

**Supplementary Table 5 |** Organic carbon and energy metabolisms and associated pathways of four major populations in SPREAD UASB microbiome (Cont.)

| Pathway and related metabolism | Name | Enzymes | EC number | Draft genome | | | |
|---|---|---|---|---|---|---|---|
| | | | | MAG.10 (Mesotoga) | MAG. 14 (Bacteroidales) | MAG.1 (Methanothrix) | MAG. 36 (Synrophales) |
| Beta-oxidation of fatty acids | fadD | Long-chain acyl-CoA synthetase | EC 6.2.1.3 | | 5 | | 1 |
| | acd | Acyl-CoA dehydrogenase | EC 1.3.8.7 | | | | 2 |
| | echA | Enoyl-CoA hydratase | EC 4.2.1.17 | | | | |
| | FadN | 3-hydroxyacyl-CoA dehydrogenase | EC 1.1.1.35 | | | | 1 |
| | fadB | 3-hydroxybutyryl-CoA dehydrogenase | EC:1.1.1.157 | | | | 2 |
| | fadA | Acetyl-CoA acyltransferase | EC 2.3.1.16 | | | | 1 |
| Fatty acid biosynthesis | fabD | [acyl-carrier-protein] S-malonyltransferase | EC:2.3.1.39 | 1 | 1 | | 2 |
| | fabF | 3-oxoacyl-[acyl-carrier-protein] synthase II | EC:2.3.1.179 | 1 | 2 | | 2 |
| | fabG | 3-oxoacyl-[acyl-carrier protein] reductase | EC:1.1.1.100 | 1 | 3 | | 3 |
| | fabH | 3-oxoacyl-[acyl-carrier-protein] synthase III | EC:2.3.1.180 | 1 | 2 | 1 | 1 |
| | fabK | Enoyl-[acyl-carrier protein] reductase II | EC:1.3.1.9 | 1 | 1 | | |
| | fabZ | 3-hydroxyacyl-[acyl-carrier-protein] dehydratase | EC:4.2.1.59 | 1 | 2 | | 1 |
| TCA cycle | acnA | Aconitate hydratase | EC 4.2.1.3 | | | 1 | 1 |
| | korA | 2-oxoglutarate/2-oxoacid ferredoxin oxidoreductase, alpha subunit | EC 1.2.7.3; EC 1.2.7.11 | 3 | 2 | 1 | 4 |
| | korB | 2-oxoglutarate/2-oxoacid ferredoxin oxidoreductase, beta subunit | EC 1.2.7.3; EC 1.2.7.11 | 3 | 2 | 1 | 4 |
| | korC | 2-oxoglutarate ferredoxin oxidoreductase subunit gamma | EC:1.2.7.3 | 2 | 1 | | 3 |
| | korD | 2-oxoglutarate ferredoxin oxidoreductase subunit delta | EC:1.2.7.3 | 2 | 1 | | 2 |
| | sdhA | Succinate dehydrogenase flavoprotein subunit | EC:1.3.5.1 | | 1 | | |
| | sdhB | Succinate dehydrogenase iron-sulfur subunit | EC:1.3.5.1 | | 1 | | |
| | sdhC | Succinate dehydrogenase cytochrome b subunit | EC:1.3.5.1 | | 1 | | |
| | sucD | Succinyl-CoA synthetase (Succinyl-CoA ligase), ADP-forming, alpha subunit | EC 6.2.1.5 | | 1 | | |
| | sucC | Succinyl-CoA synthetase (Succinyl-CoA ligase), ADP-forming, beta subunit | EC 6.2.1.5 | | 1 | | |
| | fumA | Fumarate hydratase subunit alpha | EC:4.2.1.2 | 1 | 1 | 1 | 1 |
| | fumB | Fumarate hydratase subunit beta | EC:4.2.1.2 | 1 | 1 | 1 | 1 |
| | mqo | Malate dehydrogenase (quinone or NAD-dependent) | EC 1.1.5.4; EC 1.1.1.37 | | | | |
| | pycA | Pyruvate carboxylase subunit A | EC:6.4.1.1 | | | 1 | |
| | pycB | Pyruvate carboxylase subunit B | EC:6.4.1.1 | | 1 | 1 | |

**Supplementary Table 5 |** Organic carbon and energy metabolisms and associated pathways of four major populations in SPREAD UASB microbiome (Cont.)

| Pathway and related metabolism | | Name | Enzymes | EC number | Draft genome | | | |
|---|---|---|---|---|---|---|---|---|
| | | | | | **MAG.10 (Mesotoga)** | **MAG. 14 (Bacteroidales)** | **MAG.1 (Methanothrix)** | **MAG. 36 (Synrophales)** |
| Oxidative phosphorylation chain | Complex I | nuoA | NADH-ubiquinone oxidoreductase chain A | EC 1.6.5.3 | | | | 1 |
| | | nuoB | NADH-ubiquinone oxidoreductase chain B | EC 1.6.5.3 | 1 | | | 1 |
| | | nuoC | NADH-ubiquinone oxidoreductase chain C | EC 1.6.5.3 | 1 | | | 1 |
| | | nuoD | NADH-ubiquinone oxidoreductase chain D | EC 1.6.5.3 | 1 | | | 1 |
| | | nuoE | NADH-ubiquinone oxidoreductase chain E | EC 1.6.5.3 | 2 | | | |
| | | nuoF | NADH-ubiquinone oxidoreductase chain F | EC 1.6.5.3 | 2 | | | 3 |
| | | nuoH | NADH-ubiquinone oxidoreductase chain H | EC 1.6.5.3 | 1 | | | 1 |
| | | nuoI | NADH-ubiquinone oxidoreductase chain I | EC 1.6.5.3 | | | | 1 |
| | | nuoJ | NADH-ubiquinone oxidoreductase chain J | EC 1.6.5.3 | | | | 1 |
| | | nuoK | NADH-ubiquinone oxidoreductase chain K | EC 1.6.5.3 | | | | 1 |
| | | nuoL | NADH-ubiquinone oxidoreductase chain L | EC 1.6.5.3 | | | | 1 |
| | | nuoM | NADH-ubiquinone oxidoreductase chain M | EC 1.6.5.3 | 1 | | | 1 |
| | | nuoN | NADH-ubiquinone oxidoreductase chain N | EC 1.6.5.3 | 1 | | | 1 |
| | Complex II | sdhA | Succinate dehydrogenase/fumarate reductase, flavoprotein subunit | EC 1.3.5.1 | | 1 | | |
| | | sdhB | Succinate dehydrogenase/fumarate reductase, iron/sulfur subunit | EC 1.3.5.1 | | 1 | | |
| | | sdhC | Succinate dehydrogenase/fumarate reductase, putative membrane subunit | EC 1.3.5.1 | | 1 | | |
| | Complex V (V-type) | ntpA | V-type ATP synthase subunit A | EC 3.6.3.14 | | 1 | | |
| | | ntpB | V-type ATP synthase subunit B | EC 3.6.3.14 | | 1 | | |
| | | ntpD | V-type ATP synthase subunit D | EC 3.6.3.14 | | 1 | 1 | |
| | | ntpE | V-type ATP synthase subunit E | EC 3.6.3.14 | | 1 | | |
| | | ntpI | V-type ATP synthase subunit I | EC 3.6.3.14 | | 1 | 1 | |
| | | ntpK | V-type ATP synthase subunit K | EC 3.6.3.14 | | 1 | | |
| | Complex V (F-type) | atpA | F-type H+/Na+-transporting ATPase subunit alpha | EC:7.1.2.2 | 1 | | | 2 |
| | | atpB | F-type H+-transporting ATPase subunit a | EC:7.1.2.2 | 1 | 1 | | 2 |
| | | atpC | F-type H+-transporting ATPase subunit epsilon | EC:7.1.2.2 | 1 | | | 2 |
| | | atpD | F-type H+/Na+-transporting ATPase subunit beta | EC:7.1.2.2 | 1 | | | 2 |
| | | atpF | F-type H+-transporting ATPase subunit b | EC:7.1.2.2 | 1 | | | 3 |
| | | atpE | F-type H+-transporting ATPase subunit c | EC:7.1.2.2 | 1 | 1 | | 3 |
| | | atpG | F-type H+-transporting ATPase subunit gamma | EC:7.1.2.2 | 1 | | | 2 |
| | | atpH | F-type H+-transporting ATPase subunit delta | EC:7.1.2.2 | 1 | | 1 | 1 |

**Supplementary Table 5 |** Organic carbon and energy metabolisms and associated pathways of four major populations in SPREAD UASB microbiome (Cont.)

| Pathway and related metabolism | Name | Enzymes | EC number | Draft genome | | | |
|---|---|---|---|---|---|---|---|
| | | | | **MAG.10 (Mesotoga)** | **MAG. 14 (Bacteroidales)** | **MAG.1 (Methanothrix)** | **MAG. 36 (Synrophales)** |
| Nitrogen metabolism | nrfA | Nitrite reductase (cytochrome c-552) | EC 1.7.2.2 | | 1 | | 1 |
| | nrfH | Cytochrome c nitrite reductase small subunit | | | 1 | | 1 |
| | gdhA | Glutamate dehydrogenase (NADP+) | EC 1.4.1.2; EC 1.4.1.3; EC 1.4.1.4 | 1 | 1 | 1 | 1 |
| | glt | Glutamate synthase (NAD(P)H) | EC 1.4.1.13; EC 1.4.1.14 | 2 | 2 | 1 | 1 |
| | glnA | Glutamine synthetase | EC 6.3.1.2 | 1 | 2 | 2 | 1 |
| | norB | Nitric oxide reductase subunit B | EC 1.7.2.5 | 1 | | | 1 |
| | ncd2 | Nitronate monooxygenase | EC:1.13.12.16 | 1 | 1 | | |
| Transporters | livK | Branched-chain amino acid transport system substrate-binding protein | | 4 | | 1 | 3 |
| | livH | Branched-chain amino acid transport system permease protein | | 4 | | | 3 |
| | livM | Branched-chain amino acid transport system permease protein | | 4 | | | 3 |
| | mlaD | Phospholipid/cholesterol/gamma-HCH transport system substrate-binding protein | | | 1 | | 2 |
| | mlaE | Phospholipid/cholesterol/gamma-HCH transport system permease protein | | | 1 | | 3 |
| | mlaF | Phospholipid/cholesterol/gamma-HCH transport system ATP-binding protein | | | 1 | | 2 |
| | oppA | Oligopeptide transport system substrate-binding protein | | 1 | | | |
| | oppC | Oligopeptide transport system permease protein | | 1 | | | |
| | oppD | Ligopeptide transport system ATP-binding protein | | 1 | | | |
| | oppF | Oligopeptide transport system ATP-binding protein | | 2 | | | |
| | rbsA | Ribose transport system substrate-binding protein | | 6 | | | |
| | rbsB | Ribose transport system substrate-binding protein | | 3 | | | |
| | rbsC | Ribose transport system substrate-binding protein | | 5 | | | |
| | fucp | MFS transporter, FHS family, L-fucose permease | | | 1 | | |
| | malT | Maltose/moltooligosaccharide transporter | | | 1 | | |
| | POT | Proton-dependent oligopeptide transporter, POT family | | | 1 | | |
| | znuA | Zinc transport system substrate-binding protein | | 1 | | 1 | 1 |
| | znuB | Zinc transport system permease protein | | 1 | | 1 | 1 |
| | znuC | Zinc transport system ATP-binding protein | | 1 | | 2 | 1 |

**Supplementary Table 5 |** Organic carbon and energy metabolisms and associated pathways of four major populations in SPREAD UASB microbiome (Cont.)

| **Pathway and related metabolism** | **Name** | **Enzymes** | **EC number** | **Draft genome** | | | |
|---|---|---|---|---|---|---|---|
| | | | | **MAG.10 (Mesotoga)** | **MAG. 14 (Bacteroidales)** | **MAG.1 (Methanothrix)** | **MAG. 36 (Synrophales)** |
| Transporters | pstA | Phosphate transport system permease protein | | 1 | | 1 | |
| | pstB | Phosphate transport system ATP-binding protein | | 1 | | 1 | |
| | pstC | Phosphate transport system permease protein | | 1 | | 2 | |
| | pstS | Phosphate transport system substrate-binding protein | | | | 5 | |
| | feoA | Ferrous iron transport protein A | | 1 | | 3 | |
| | feoB | Ferrous iron transport protein B | | 1 | 1 | 1 | 1 |
| | lptB | Lipopolysaccharide export system ATP-binding protein | | 1 | 1 | | 1 |
| | lptF | Lipopolysaccharide export system permease protein | | | 1 | | 1 |
| | lptG | Lipopolysaccharide export system permease protein | | | | | 1 |
| | lolA | Outer membrane lipoprotein carrier protein | | | 1 | | 1 |
| | lolC | Lipoprotein-releasing system permease protein | | 1 | 2 | 2 | 1 |
| | lolD | Lipoprotein-releasing system ATP-binding protein | | 1 | 1 | 1 | 1 |
| | wtpA | Molybdate/tungstate transport system substrate-binding protein | | | 1 | | |
| | wtpB | Molybdate/tungstate transport system permease protein | | | 1 | | |
| | wtpC | Molybdate/tungstate transport system ATP-binding protein | | | | 1 | |
| | fdhC | Formate transporter | | 1 | 1 | | |
| | ugpA | Sn-glycerol 3-phosphate transport system permease protein | | 2 | | | |
| | ugpB | Sn-glycerol 3-phosphate transport system substrate-binding protein | | 5 | | | |
| | ugpE | Sn-glycerol 3-phosphate transport system permease protein | | 2 | | | |
| | art | Arginine/lysine/histidine transport system permease protein | | 3 | | | |
| | thuE | Trehalose/maltose transport system substrate-binding protein | | 1 | | | |
| | thuF | Trehalose/maltose transport system permease protein | | 1 | | | |
| | thuG | Trehalose/maltose transport system permease protein | | 1 | | | |
| | malE | Maltose/maltodextrin transport system substrate-binding protein | | 3 | | | |
| | malF | Maltose/maltodextrin transport system permease protein | | 2 | | | |
| | malG | Maltose/maltodextrin transport system permease protein | | 1 | | | |
| | ganQ | Arabinogalactan oligomer / maltooligosaccharide transport system permease protein | | 3 | | | |
| | msmE | Raffinose/stachyose/melibiose transport system substrate-binding protein | | 1 | | | |
| | msmF | Raffinose/stachyose/melibiose transport system permease protein | | 2 | | | |
| | msmG | Raffinose/stachyose/melibiose transport system permease protein | | 1 | | | |

**Supplementary Table 5 |** Organic carbon and energy metabolisms and associated pathways of four major populations in SPREAD UASB microbiome (Cont.)

| **Pathway and related metabolism** | | **Name** | **Enzymes** | **EC number** | **Draft genome** | | | |
|---|---|---|---|---|---|---|---|---|
| | | | | | **MAG.10 (Mesotoga)** | **MAG. 14 (Bacteroidales)** | **MAG.1 (Methanothrix)** | **MAG. 36 (Synrophales)** |
| Transporters | | msmX | Multiple sugar transport system ATP-binding protein | | 1 | | | |
| | | chiF | Putative chitobiose transport system permease protein | | 2 | | | |
| | | chiG | Putative chitobiose transport system permease protein | | 1 | | | |
| | | gtsA | Glucose/mannose transport system substrate-binding protein | | 1 | | | |
| | | gtsB | Glucose/mannose transport system permease protein | | 1 | | | |
| | | gtsC | Glucose/mannose transport system permease protein | | 1 | | | |
| | | ABC. MS.P | Multiple sugar transport system permease protein | | 12 | | | |
| | | ABC. MS.P1 | Multiple sugar transport system permease protein | | 14 | | | |
| | | ABC. MS.S | Multiple sugar transport system substrate-binding protein | | 13 | | | |
| | | ABC.S S.P | Simple sugar transport system permease protein | | 2 | | | |
| | | ABC.S S.A | Simple sugar transport system ATP-binding protein | | 1 | | | |
| Stress response | Multiple drugs | acrA | Membrane fusion protein, multidrug efflux system | | | 1 | | 3 |
| | | oprM | Outer membrane protein, multidrug efflux system | | | | | 1 |
| | | mdtG | Membrane fusion protein, multidrug efflux system | | | | | 1 |
| | | mdtK | MATE family, multidrug efflux pump | | 1 | 1 | | |
| | Heavy metal | cusA | Cu(I)/Ag(I) efflux system membrane protein CusA/SilA | | | 1 | | |
| | | cusB | Membrane fusion protein, Cu(I)/Ag(I) efflux system | | | 1 | | 1 |
| | | czcC | Outer membrane protein, cobalt-zinc-cadmium efflux system | | | | | 1 |
| | | arsC | Arsenate reductase (thioredoxin) | | | 1 | 2 | |
| | | arsB | Arsenite transporter | | 1 | 1 | | 1 |
| Secretion system | Sec-SRP | ftsY | Fused signal recognition particle receptor | | 1 | 1 | 1 | 1 |
| | | secA | Preprotein translocase subunit SecA | | 1 | 1 | | 1 |
| | | secE | Preprotein translocase subunit SecE | | 1 | 1 | 1 | 1 |
| | | secD | Preprotein translocase subunit SecD | | 1 | | 1 | 1 |
| | | secF | Preprotein translocase subunit SecF | | 1 | | 1 | 1 |
| | | secG | Preprotein translocase subunit SecG | | 1 | 1 | | 1 |
| | | secY | Preprotein translocase subunit SecY | | 1 | 1 | 1 | 1 |
| | | yajC | Preprotein translocase subunit YajC | | 1 | 1 | | 1 |
| | | yidC | YidC/Oxa1 family membrane protein insertase | | 1 | 1 | | 1 |
| | | ffh | Signal recognition particle subunit SRP54 | EC 3.6.5.4 | 1 | 1 | 1 | 1 |
| | Twin arginine targeting | tatA | Sec-independent protein translocase protein TatA | | | 1 | | 2 |
| | | tatC | Sec-independent protein translocase protein TatC | | | 1 | | 1 |

**Supplementary Table 5 |** Organic carbon and energy metabolisms and associated pathways of four major populations in SPREAD UASB microbiome (Cont.)

| **Pathway and related metabolism** | | **Name** | **Enzymes** | **EC number** | **Draft genome** | | | |
|---|---|---|---|---|---|---|---|---|
| | | | | | **MAG.10 (Mesotoga)** | **MAG. 14 (Bacteroidales)** | **MAG.1 (Methanothrix)** | **MAG. 36 (Synrophales)** |
| Carbohydrate-degradation | Cellulases | GH5 | Cellulase | EC3.2.1.4 | 1 | 1 | | |
| | | GH9 | Endoglucanase | EC3.2.1.4 | | 1 | | |
| | Beta-glucosidases/ beta-xylosidases | GH1 | Beta-glucosidase | EC3.2.1.21 | 2 | | | |
| | | GH3 | β-glucosidase | EC3.2.1.21 | 3 | | 1 | |
| | Hemicellulose | GH10 | Endo-1,4-beta-xylanase | EC 3.2.1.8 | 1 | | | |
| | Beta-glucanases | GH16 | β-glucosidase | EC 3.2.1.21 | | 1 | | |
| | | GH144 | Endo-β-1,2-glucanase | EC3.2.1.71 | 2 | | | |
| | Alpha-glucanases | GH13 | α-amylase | EC3.2.1.1 | 8 | 6 | 3 | 1 |
| | | GH23 | Peptidoglycanases | EC3.2.1.17 | | 3 | 5 | |
| | | GH31 | α-glucosidase | EC3.2.1.20 | 1 | | | |
| | | GH57 | α-amylase | EC 3.2.1.1 | 1 | 1 | 2 | 1 |
| | | GH77 | Amylomaltase or 4-α-glucanotransferase | EC2.4.1.25 | | 1 | 1 | |
| | Chitinases | GH18 | Chitinase | EC3.2.1.14 | | 1 | | |
| | | GH20 | N-acetyl β-glucosaminidase | EC3.2.1.52 | | 4 | | |
| | Peptidoglycanases | GH23 | Peptidoglycanases | EC3.2.1.17 | | 3 | 5 | |
| | Other hemicellulases | GH35 | β-galactosidase | EC3.2.1.23 | 2 | | | |
| | | GH36 | α-galactosidase | EC3.2.1.22 | 1 | | | |
| | | GH51 | Xylan β-1,4-xylosidase | EC 3.2.1.37 | 2 | | | |
| | | GH97 | α-galactosidase | EC3.2.1.22 | | 1 | | |
| | Glycoconjugate-degrading enzymes | GH38 | α-mannosidase | EC3.2.1.24 | 1 | | | |
| | | GH84 | Hyaluronidase | EC3.2.1.35 | 1 | | | |
| | | GH92 | α-mannosidase | EC3.2.1.24 | | 2 | | |
| | | GH109 | α-N-acetylgalactosaminidase | EC3.2.1.49 | 4 | | | |
| | Pectinases | GH127 | β-L-arabinofuranosidase | EC 3.2.1.185 | 2 | | | |
| | | GH142 | β-L-arabinofuranosidase | EC3.2.1.185 | 1 | | | |
| | Mannanases | GH130 | β-1,2-mannosidase | EC 3.2.1.197 | 3 | | | |
| Protease | | C01A | Peptidase family C1 contains many endopeptidases and a few exopeptidases | | | 1 | | |
| | | C01B | Peptidase family C1 contains many endopeptidases and a few exopeptidases | | | 2 | | |
| | | C69 | Peptidase family C69 contains dipeptidases and aminopeptidases | | 7 | 3 | | |

**Supplementary Table 5 |** Organic carbon and energy metabolisms and associated pathways of four major populations in SPREAD UASB microbiome (Cont.)

| **Pathway and related metabolism** | | **Name** | **Enzymes** | **EC number** | **Draft genome** | | | |
|---|---|---|---|---|---|---|---|---|
| | | | | | **MAG.10 (Mesotoga)** | **MAG. 14 (Bacteroidales)** | **MAG.1 (Methanothrix)** | **MAG. 36 (Synrophales)** |
| Protease | | M03A | Peptidase family M3 contains metallopeptidases with varied activities | | | 2 | | |
| | | M20A | Peptidase family M20 contains exopeptidases: carboxypeptidases, dipeptidases and a specialised aminopeptidase | | 3 | 1 | | |
| | | M20B | | | 1 | 2 | | |
| | | M20C | | | | 2 | 1 | |
| | | M20D | | | 2 | | | |
| | | M20F | | | | 1 | | |
| | | M20X | | | | 1 | | 1 |
| | | M32 | Peptidase family M32 contains metallocarboxypeptidases | | 1 | | | |
| | | M42 | Peptidase family M42 contains metalloaminopeptidases some of which also have acylaminoacylpeptidase activity | | 5 | | | |
| | | M55 | Peptidase family M55 contains an aminopeptidase and a number of uncharacterised putative peptidases | | 1 | 1 | | |
| | | S09B | Peptidase family S9 contains a varied set of serine-dependent peptidases. | | | 1 | | |
| | | S37 | Peptidase family S37 contains a tripeptidyl-peptidase | | | 1 | | |
| | | S46 | Peptidase family S46 contains dipeptidyl-peptidases | | | 2 | | |
| | | T02 | Peptidase family T2 contains a set of the N-terminal nucleophile hydrolases | | 1 | 1 | | |
| | | T03 | Peptidase family T3 contains self-processing proteins that express aminopeptidase | | 3 | | | |
| Motility system | Flagellar system | flgA | Flagellar basal body P-ring formation protein FlgA | | | | | 1 |
| | | flgB | Flagellar basal-body rod protein FlgB | | | | | 1 |
| | | flgC | Flagellar basal-body rod protein FlgC | | | | | 1 |
| | | flgD | Flagellar basal-body rod modification protein FlgD | | | | | 1 |
| | | flgH | Flagellar L-ring protein FlgH | | | | | 1 |
| | | flgI | Flagellar P-ring protein FlgI | | | | | 1 |
| | | flgK | Flagellar hook-associated protein 1 | | | | | 1 |
| | | flgM | Negative regulator of flagellin synthesis | | | | | 1 |
| | | flgR | Two-component system, NtrC family, response regulator | | | | | 1 |
| | | fliA | RNA polymerase sigma factor for flagellar operon | | | | | 1 |
| | | fliC | Flagellin | | | | | 1 |
| | | fliE | Flagellar hook-basal body complex protein | | | | | 1 |
| | | fliI | Flagellum-specific ATP synthase | EC 7.4.2.8 | | | | 1 |
| | | fliJ | Flagellar protein FliJ | | | | | 1 |
| | | motB | Chemotaxis protein | | | | | 3 |

**Supplementary Table 5 |** Organic carbon and energy metabolisms and associated pathways of four major populations in SPREAD UASB microbiome (Cont.)

| Pathway and related metabolism | | Name | Enzymes | EC number | Draft genome | | | |
|---|---|---|---|---|---|---|---|---|
| | | | | | MAG.10 (Mesotoga) | MAG. 14 (Bacteroidales) | MAG.1 (Methanothrix) | MAG. 36 (Synrophales) |
| Motility system | Pilus system | pilB | Type IV pilus assembly protein | | 1 | | | 1 |
| | | pilC | Type IV pilus assembly protein | | 1 | | | |
| | | pilO | Type IV pilus assembly protein | | | | | 1 |
| | | pilP | Type IV pilus assembly protein | | | | | 1 |
| | | pilQ | Type IV pilus assembly protein | | | | | 1 |
| | | pilR | Type IV pilus assembly protein | | | | | 1 |
| | | pilN | Type IV pilus assembly protein | | | | | 1 |
| | | pilM | Type IV pilus assembly protein | | | | | 1 |
| | | pilT | Twitching motility protein | | 1 | | | 3 |
| | | pilY1 | Type IV pilus assembly protein | | | | | 1 |
| Methanogenesis | | acs | Acetyl-CoA synthetase | EC:6.2.1.1 | | | 4 | |
| | | fwdA | Formylmethanofuran dehydrogenase subunit A | EC:1.2.7.12 | | | 2 | |
| | | fwdB | Formylmethanofuran dehydrogenase subunit B | EC:1.2.7.12 | | | 3 | |
| | | fwdC | Formylmethanofuran dehydrogenase subunit C | EC:1.2.7.12 | | | 2 | |
| | | fwdD | Formylmethanofuran dehydrogenase subunit D | EC:1.2.7.12 | | | 3 | |
| | | fwdE | Formylmethanofuran dehydrogenase subunit E | EC:1.2.7.12 | | | 2 | 1 |
| | | fwdF | 4Fe-4S ferredoxin | EC:1.2.7.12 | | | 2 | |
| | | fwdG | 4Fe-4S ferredoxin | EC:1.2.7.12 | | | 1 | |
| | | ftr | Formylmethanofuran--tetrahydromethanopterin N-formyltransferase | EC:2.3.1.101 | | | 1 | |
| | | mch | Methenyltetrahydromethanopterin cyclohydrolase | EC:3.5.4.27 | | | 1 | |
| | | mtd | Methylenetetrahydromethanopterin dehydrogenase | EC:1.5.98.1 | | | 1 | |
| | | mer | 5,10-methylenetetrahydromethanopterin reductase | EC:1.5.98.2 | | | 1 | |
| | | mtrA | Tetrahydromethanopterin S-methyltransferase subunit A | EC:7.2.1.4 | | | 2 | |
| | | mtrB | Tetrahydromethanopterin S-methyltransferase subunit B | EC:7.2.1.4 | | | 1 | |
| | | mtrC | Tetrahydromethanopterin S-methyltransferase subunit C | EC:7.2.1.4 | | | 1 | |
| | | mtrD | Tetrahydromethanopterin S-methyltransferase subunit D | EC:7.2.1.4 | | | 1 | |
| | | mtrE | Tetrahydromethanopterin S-methyltransferase subunit E | EC:7.2.1.4 | | | 1 | |
| | | mtrF | Tetrahydromethanopterin S-methyltransferase subunit F | EC:7.2.1.4 | | | 1 | |
| | | mtrG | Tetrahydromethanopterin S-methyltransferase subunit G | EC:7.2.1.4 | | | 1 | |
| | | mtrH | Tetrahydromethanopterin S-methyltransferase subunit H | EC:7.2.1.4 | | | 2 | |
| | | mcrA | Methyl-coenzyme M reductase alpha subunit | EC:2.8.4.1 | | | 1 | |
| | | mcrB | Methyl-coenzyme M reductase beta subunit | EC:2.8.4.1 | | | 1 | |
| | | mcrC | Methyl-coenzyme M reductase subunit C | EC:2.8.4.1 | | | 1 | |
| | | mcrD | Methyl-coenzyme M reductase subunit D | EC:2.8.4.1 | | | 1 | |
| | | mcrG | Methyl-coenzyme M reductase gamma subunit | EC:2.8.4.1 | | | 1 | |

**Supplementary Table 6 |** Accession numbers of previously published sequencing data for source tracking and PCoA analysis in this study.

| Accession number | Source types | Samples | Hypervariable | Platform |
|---|---|---|---|---|
| DRP002796 | UASB | 6 | V4 | Illumina MiSeq |
| DRP003910 | UASB | 42 | V4 | Illumina MiSeq |
| DRP003910 | UASB | 5 | V4 | Illumina MiSeq |
| PRJNA522831 | UASB | 5 | V4 | Illumina MiSeq |
| PRJNA522972 | UASB | 34 | v4 | Illumina MiSeq |
| DRP003399 | Anaerobic digestion | 139 | V4 | Illumina MiSeq |
| PRJNA315957 | Anaerobic digestion | 66 | V3-V4 | 454 GS FLX Titanium |
| PRJNA328964 | Anaerobic digestion | 20 | V4 | Illumina MiSeq |
| PRJNA507417 | Anaerobic digestion | 52 | V4 | Illumina HiSeq |
| PRJEB5095 | Activate sludge | 44 | V4 | Illumina HiSeq |
| PRJNA306487 | Activate sludge | 12 | V4 | GS Junior Titanuim |
| PRJNA448640 | Activate sludge | 100 | V4 | Illumina HiSeq |

**Supplementary Table 7 |** Life cycle inventory of CAD, TAAD and SPREAD

| Process | Type | Flow | Unit | CAD | TAAD | SPREAD |
|---|---|---|---|---|---|---|
| Pretreatment | In | Thickened sludge (97% moisture content) | t | / | 1 | 1 |
| | In | Heat (completely provided by biogas combustion) | GJ | / | 0.13 | 0.13 |
| | In | NaOH | kg | / | 5.82 | 5.82 |
| | In | HCl | kg | / | 5.23 | 5.23 |
| | In | Electricity (Pretreatment) | kWh | / | 10 | 10 |
| | Out | Thermal-alkaline sludge (97% moisture content) | t | / | 1 | 1 |
| | Out | liquefied sludge (99.9% moisture content) | t | / | / | 0.75 |
| | Out | Ash layer (25.2kg dry solid and 222.53kg water) | t | / | / | 0.2477 |
| Anaerobic digestion (with cogeneration) | In | Thickened sludge (97% moisture content) | t | 1 | / | / |
| | In | Thermal-alkaline sludge (97% moisture content) | t | / | 1 | / |
| | In | liquefied sludge (99.9% moisture content) | t | / | / | 0.75 |
| | In | Electricity (anaerobic digester) | kWh | 2.5 | 2.5 | 0.25 |
| | In | Heat (anaerobic digester, provided by biogas combustion) | GJ | 0.13 | / | / |
| | In | Electricity (compressors, removal the moisture and $H_2S$ in biogas) | kWh | 0.0625 | 0.0625 | 0.0625 |
| | Out | Emission to air, biogenic $CH_4$ (biogas loss) | kg | 0.0928 | 0.1326 | 0.0353 |
| | Out | Emission to air, biogenic $CO_2$ (biogas loss) | kg | 0.1708 | 0.2440 | 0.0650 |
| | Out | Emission to air, CO (emission from the combustion of biogas) | g | 49.1040 | 63.8352 | 56.1150 |
| | Out | Emission to air, $NH_3$ (biogas loss) | g | 1.7061 | 2.8420 | 1.8569 |
| | Out | Emission to air, $N_2O$ (emission from the combustion of biogas) | g | 0.2520 | 0.3276 | 0.2880 |
| | Out | Emission to air, $NO_x$ (emission from the combustion of biogas) | g | 32.0400 | 41.6520 | 36.6146 |
| | Out | Emission to air, $CH_4$ (emission from the combustion of biogas) | g | 68.7600 | 89.3880 | 78.5774 |
| | Out | Anaerobically-digested sludge (27.575kg dry solid and 0.97t water) | t | 0.9976 | / | / |
| | Out | Anaerobically-digested sludge (24.495kg dry solid and 0.97t water) | t | / | 0.9945 | / |
| | Out | Wastewater | t | / | / | 0.7477 |
| Filter pressing dewatering | In | Anaerobically-digested sludge (27.575kg dry solid and 0.97t water) | t | 0.9976 | / | / |
| | In | Anaerobically-digested sludge (24.495kg dry solid and 0.97t water) | t | / | 0.9945 | / |
| | In | Ash layer (25.2kg dry solid and 222.53kg water) | t | / | / | 0.2477 |
| | In | Electricity | kWh | 1.2475 | 1.2350 | 0.3100 |
| | In | $FeCl_3$ | kg | 4.1575 | 3.9500 | 1.0310 |
| | In | PAM | kg | 3.7000 | 3.3100 | 0.9180 |
| | Out | Digestate (70% moisture content, 27.575kg dry solid and 64.342kg water) | t | 0.0919 | / | / |
| | Out | Digestate (70% moisture content, 24.495kg dry solid and 57.155kg water) | t | / | 0.0817 | / |
| | Out | Digestate (70% moisture content, 25.2kg dry solid and 58.8kg water) | t | / | / | 0.0840 |

| Process | Type | Flow | Unit | CAD | TAAD | SPREAD |
|---|---|---|---|---|---|---|
| | Out | Wastewater | t | 0.9057 | 0.9128 | 0.1637 |
| Incineration | In | Digestate (70% moisture content, 27.575kg dry solid and 64.342kg water) | t | 0.0919 | / | / |
| | In | Digestate (70% moisture content, 24.495kg dry solid and 57.155kg water) | t | / | 0.0817 | / |
| | In | Digestate (70% moisture content, 25.2kg dry solid and 58.8kg water) | t | / | / | 0.0840 |
| | In | Electricity | kWh | 0.4750 | 0.4750 | 0.4750 |
| | In | Heat (produced by diesel) | MJ | 0.9875 | 0.9875 | 0.9875 |
| | In | Heavy fuel (burned in furnace) | kg | 0.8538 | 0.7583 | 0.7803 |
| | In | NaOH | kg | 0.3360 | 0.2984 | 0.3071 |
| | In | Lime (hydrated, loose) | kg | 0.1366 | 0.1213 | 0.1248 |
| | In | Ammonia (liquid) | kg | 0.1025 | 0.0910 | 0.0936 |
| | Out | Emission to air, $CO_2$ | kg | 4.3065 | 3.8251 | 3.9360 |
| | Out | Emission to air, CO | mg | 0.0041 | 0.0037 | 0.0038 |
| | Out | Emission to air, $NO_2$ | mg | 0.0275 | 0.0245 | 0.0252 |
| | Out | Emission to air, Particulates, < 10um ($PM_{10}$) | µg | 0.0551 | 0.0489 | 0.0503 |
| | Out | Emission to air, Furans | pg | 0.0014 | 0.0012 | 0.001493 |
| | Out | Ash | kg | 23.2630 | 20.6650 | 21.2590 |
| Landfill | In | Ash | kg | 23.2630 | 20.6650 | 21.2590 |
| | Out | Emission to water, $PO_4^{3-}$ | kg | 0.0080 | 0.0071 | 0.0073 |
| | Out | Emission to water, $NO_3^-$ | kg | 0.1695 | 0.1505 | 0.1549 |
| | Out | Emission to soil, total arsenic (T-As) | g | 0.2110 | 0.1874 | 0.1928 |
| | Out | Emission to soil, total cadmium (T-Cd) | g | 0.0080 | 0.0071 | 0.0073 |
| | Out | Emission to soil, total chromium (T-Cr) | g | 0.8390 | 0.7450 | 0.7667 |
| | Out | Emission to soil, total copper (T-Cu) | g | 1.1958 | 1.0619 | 1.0928 |
| | Out | Emission to soil, total lead (T-Pb) | g | 0.3290 | 0.2922 | 0.3007 |
| | Out | Emission to soil, total mercury (T-Hg) | g | 0.0016 | 0.0014 | 0.0015 |
| | Out | Emission to soil, total nickel (T-Ni) | g | 0.2680 | 0.2380 | 0.2449 |
| | Out | Emission to soil, total Zinc (T-Zn) | g | 3.9050 | 3.4676 | 3.5687 |
| Nutrients recovery | In | Wastewater | t | 0.9057 | 0.9128 | 0.9115 |
| | In | Electricity (Nutrients recovery) | kWh | 0.2505 | 0.2536 | 0.2535 |
| | In | $FeCl_3$ | kg | 0.5400 | 0.5440 | / |
| | In | $MgCl_2$ | kg | / | / | 4.6870 |
| | Out | Emission to air, NO | g | 0.1510 | 0.1520 | 0.0500 |
| | Out | Wastewater | t | 0.9057 | 0.9128 | 0.9115 |

| Process | Type | Flow | Unit | CAD | TAAD | SPREAD |
|---|---|---|---|---|---|---|
| Avoided products | Out | Electricity (recovered from combining heat and power (CHP) system, in which 35% of energy in biogas (23 MJ/$m^3$) is converted to electricity and 50% is converted to thermal energy in hotwater at 90 ºC) | kWh | 16.1000 | 23.0000 | 18.3987 |
| | Out | Protein feed, 100% crude | kg | / | / | 2.27 |
| | Out | Inorganic phosphorus fertilizer | kg | / | / | 2.26 |
| Transportation to disposal sites | - | 20-ton diesel lorry | tkm | 1.1632 | 1.0333 | 1.0630 |

Note: LCI data were mainly derived from the survey and experimental data (Nielsen et al., 2010; Yoshida et al., 2014; Schaubroeck et al., 2015; Li et al., 2017; Yoshida et al., 2018; Zhou et al., 2022); background inventory data including the electricity, chemicals, and materials used in this study were obtained from the Ecoinvent 3.5 database.
